# PARprolink: a photoaffinity probe for identifying poly(ADP-ribose)-binding proteins

**DOI:** 10.1101/2020.12.28.424596

**Authors:** Morgan Dasovich, Morgan Q. Beckett, Scott Bailey, Shao-En Ong, Marc M. Greenberg, Anthony K. L. Leung

**Affiliations:** Department of Chemistry, Krieger School of Arts and Sciences, Johns Hopkins University, Baltimore, MD 21218, USA; Department of Biochemistry and Molecular Biology, Bloomberg School of Public Health, Johns Hopkins University, Baltimore, MD 21205, USA; Department of Biophysics and Biophysical Chemistry, School of Medicine, Johns Hopkins University, Baltimore, MD 21205, USA; Department of Pharmacology, University of Washington, Seattle, WA 98195, USA; Department of Molecular Biology and Genetics and Department of Oncology, School of Medicine, Johns Hopkins University, Baltimore, MD 21205, USA

## Abstract

Post-translational modification of proteins with poly(ADP-ribose) (PAR) is an important component of the DNA damage response. Four PAR synthesis inhibitors have recently been approved for the treatment of breast, ovarian, and prostate cancers. Despite its clinical significance, a molecular understanding of PAR function, including its binding partners, remains incomplete. In this work, we synthesize a PAR photoaffinity probe that captures and isolates endogenous PAR binders. Our method identified dozens of known PAR-binding proteins and hundreds of novel binders involved in DNA repair, RNA processing, and metabolism. PAR binding by eight candidates was confirmed using pull-down and/or electrophoretic mobility shift assays. Using PAR probes of defined lengths, we detected proteins that preferentially bind to 40-mer over 8-mer PAR, indicating that polymer length may regulate the outcome and timing of PAR signaling pathways. This investigation produces the first census of PAR-binding proteins, provides a proteome-wide view of length-selective PAR binding, and associates PAR binding with RNA metabolism and the formation of biomolecular condensates.

---

Poly(ADP-ribose) (PAR) is an NAD^+^-dependent post-translational modification (PARylation) synthesized by PAR polymerases (PARPs).^1–6^ PARP inhibitors, which have garnered four FDA approvals in the past 6 years, are used to treat cancers.^7^ Preclinical data also support repurposing these anticancer drugs as therapeutics for neurodegeneration, cardiac failure and inflammation.^8^ A major function of PARylation is the recruitment of proteins through non-covalent interactions. For instance, PARP1 synthesizes PAR within seconds following DNA strand scission.^9^ These PAR chains in turn recruit DNA repair proteins to the damaged sites, locally concentrating repair processes precisely where and when they are needed on the chromatin.^10^ Understanding of PAR-dependent processes has been bolstered by proteomics methods that have identified thousands of ADP-ribosylated proteins during genotoxic stress and other contexts.^10–16^ However, complementary methods that identify non-covalent PAR interactions are currently lacking. Here, we describe a photo-crosslinking-based proteomics approach that identifies PAR-binding proteins.

Antibody-based approaches have been used to characterize the PAR interactome, which includes PAR binders, PARylated proteins and indirect interactors, making it difficult to identify direct PAR-protein interactions.^17,18^ To date, 92 proteins have been shown to bind PAR directly.^19,20^ This number is relatively small compared to RNA- and DNA-binding proteins (1,541 and 2,765 respectively).^21,22^ Therefore, we reasoned that many PAR-binding proteins remain undiscovered. A census of the PAR-binding proteome would provide greater insight into PAR-dependent pathways, such as DNA repair, and may reveal novel biology.

Photo-cross-linking strategies have been used widely to identify proteins that bind to RNA.^23,24^ Cross-linking is advantageous since it covalently traps binders, allowing for stringent washes that remove indirect interactions. Since PAR and RNA are structurally similar, we envisioned employing a cross-linking strategy to identify PAR-binding proteins. We synthesized a photoaffinity probe (PARprolink, Figure 1a) consisting of: PAR of defined length, a biotin handle for enrichment, and a single, randomly incorporated photo-inducible cross-linker to stabilize PAR-protein interactions. PARs of defined-length were purified from an *in vitro* enzymatic reaction using anion exchange chromatography.^25^ The biotin handle was incorporated at the 2’-OH-terminus of PAR using the ELTA bioconjugation technique.^26^ We took advantage of the selective modification of RNA hydroxyl groups by activated carboxylic acids to randomly incorporate the benzophenone tethered photo-inducible cross-linker on PAR via a nicotinic acid imidazolide (R).^27–29^ Polymers containing a single benzophenone modification were purified from a PAR mixture composed mostly of 0, 1 and 2 conjugated nicotinic acid analogues by C_18_-reverse phase HPLC (Figure 1b and S1a).

**Figure 1.**
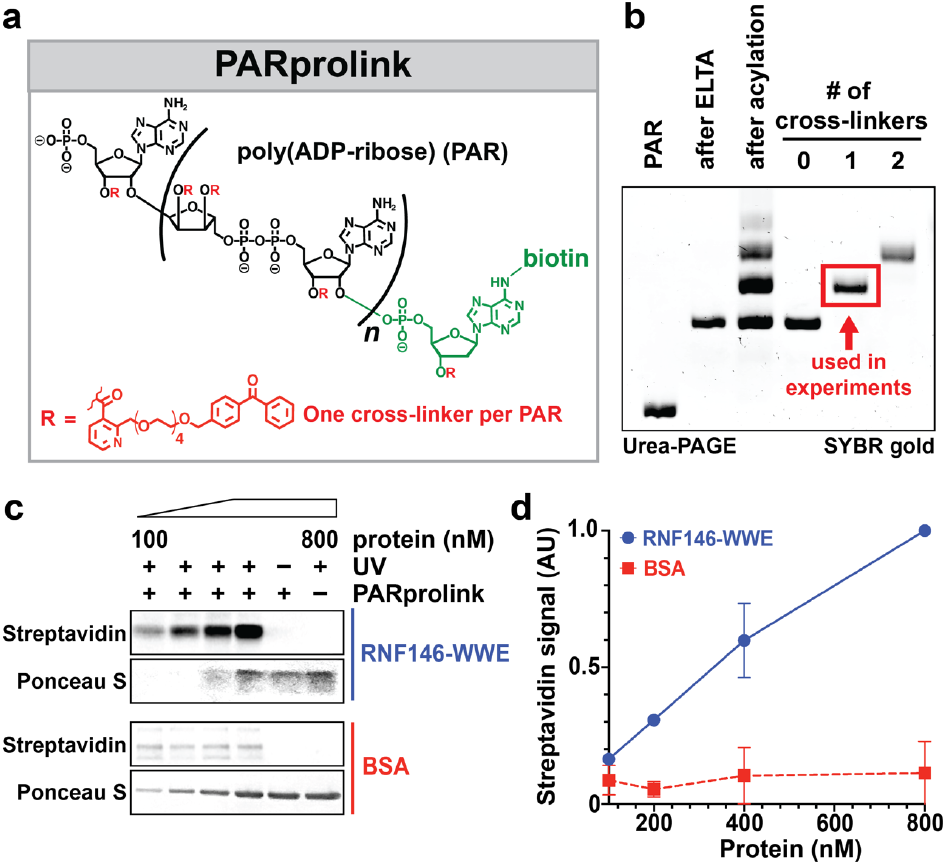
PARprolink selectively captures PAR-binding proteins *in vitro*. (a) PARprolink structure, where each PAR molecule contains one randomly incorporated cross-linker. (b) PAGE analyses of PARprolink at each synthesis step. (c) PARprolink (100 nM) was mixed with the indicated concentrations of protein, irradiated at 350 nm, and PAR-protein cross-link formation detected with streptavidin. (d) Quantification of streptavidin signal from c, values represent mean ± s.d. (*n* = 3).

The specificity of PARprolink for PAR-binding proteins was examined by incubating the probe with an increasing amount of either the PAR-binding WWE domain from human RNF146 or bovine serum albumin (BSA). Mixtures were then irradiated (350 nm, 10 min), separated by SDS-PAGE and transferred to nitrocellulose, which selectively retains protein but not PAR.^30^ Irradiation of PARprolink incubated with WWE, but not BSA, formed a protein cross-link that could be detected with streptavidin in a dose-dependent manner (Figure 1c and S1b). This signal was dependent on UV irradiation and the presence of PARprolink, consistent with covalent conjugation of the PAR-binding domain to the biotinylated probe. PARprolink specificity was further demonstrated in a complex background by cross-linking WWE domain dosed in cell extracts (Figure S1c). Addition of unlabeled PAR reduced the streptavidin signal on WWE (Figure S1d), suggesting that cross-link formation depends on the PAR-WWE interaction.

To survey the human PAR-binding proteome, we irradiated HeLa nuclear extract incubated with either an 8-mer PARprolink or biotinylated 8-mer PAR lacking benzophenone (negative control). PAR-protein cross-links were isolated with streptavidin, then subjected to on-bead trypsin digestion (Figure 2a, Figure S2a; Supplementary Data File). LC-MS/MS analysis of the pull-downs identified 798 proteins with two unique peptides in two replicates, and their abundance was quantified using the label-free quantification technology MaxLFQ (Figure S2b).^31^ The majority (743, 93%) were at least two-fold more abundant in the pull-down with PARprolink than in the no-cross-linker control (Figure 2b), demonstrating that the stringent wash removed most non-covalent interactions. There was no correlation between LFQ intensity and protein copy number, further indicating a specific enrichment by PARprolink (Figure S2c). The identified proteins overlap significantly with those identified by two antibody-based PAR interactome studies (*P* = 6.05 x 10^-111^, 1.10 x 10^-103^, Figure S2d and Supplementary Datafile).^17,18^ Although PARprolink identified a similar number of overall proteins as either antibody-based study, our cross-linking approach captured a greater percentage of known PAR binders (39 out of 92 (42%; Figure S2e, Supplementary Datafile).^19,20^

**Figure 2.**
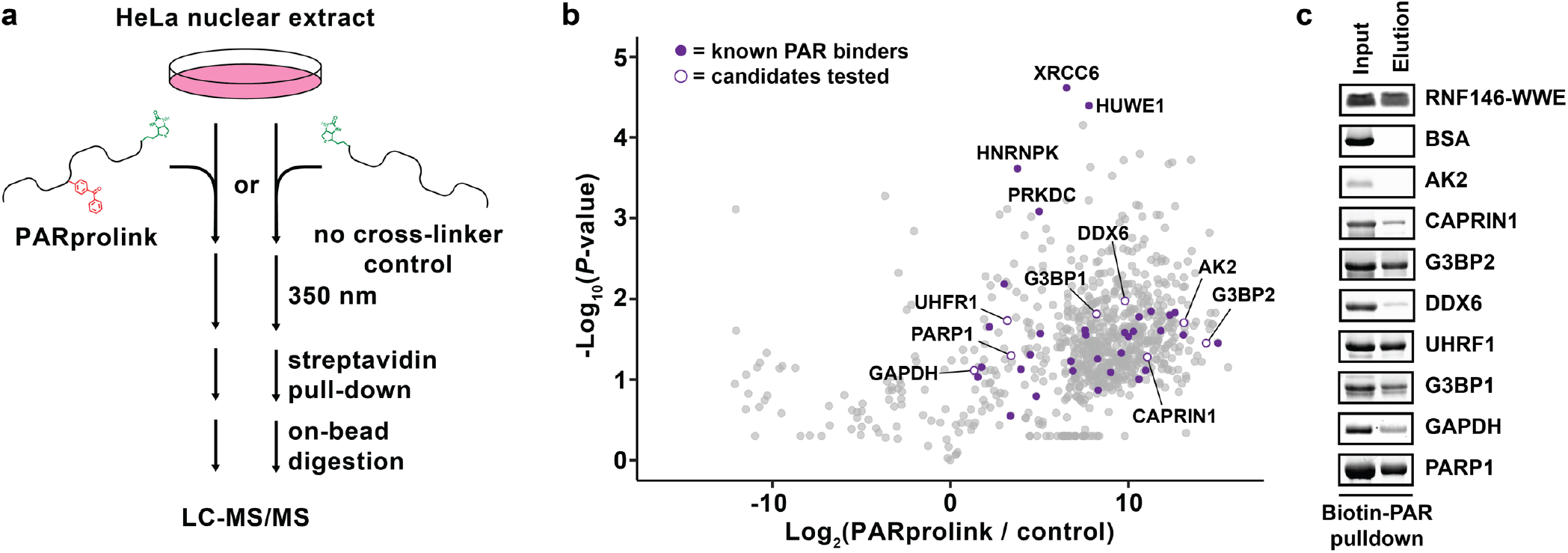
Photo-affinity-based isolation of the endogenous PAR-binding proteome. (a) Proteomics workflow schematic. (b) Volcano plot of protein enrichment ratios and -Log10(*P*-values) from proteomics experiments (*n* = 2). (c) Biotin-PAR pull-downs with candidate PARbinding proteins.

To further validate that PARprolink identifies direct PAR-protein interactions, we expressed and purified eight candidates (AK2, CAPRIN1, DDX6, G3BP2, UHRF1, G3BP1, GAPDH, PARP1) and subjected them to two PAR-binding assays. Initially, the candidate proteins were incubated with a mixture of biotinylated PAR of different lengths, followed by streptavidin pull-down. The specificity of the assay was validated with the WWE domain and BSA as a negative control. We confirmed a direct interaction between PAR and seven out of eight candidates (Figure 2c and S3).

The affinities of these candidates for 16-mer PAR were determined by electromobility shift assays (EMSAs) (Table 1, Figure S4). Consistent with our qualitative pull-down assay, the same seven candidates had affinities for PAR within the range reported for other PAR-binding proteins (*K_D_* = 1 nM-10 μM).^20,32^ The dissociation constant for AK2, which was not detected in the pull-down experiment, was estimated to be 33-65 μM, suggesting the AK2-PAR interaction may not be physiologically relevant. Taken together, these results indicate that PARprolink captured a substantial fraction of the known PAR-binding proteome and enabled the discovery of novel PAR binders.

**Table 1.** Summary of PAR-protein affinities measured with EMSAs

| Protein | $K_D$ (nM) | 95% C.I. (nM) | $R^2$ |
| --- | --- | --- | --- |
| AK2 | ~46,000 | 33,000-65,000 | 0.88 |
| CAPRIN1 | ~3,400 | 1,500-7,700 | 0.82 |
| G3BP2 | ~313 | 197-497 | 0.91 |
| DDX6 | ~199 | 134-291 | 0.94 |
| UHRF1 | ~162 | 115-230 | 0.92 |
| G3BP1 | ~93 | 73-124 | 0.95 |
| GAPDH | ~54 | 37-79 | 0.94 |
| PARP1 | ~27 | 20-38 | 0.93 |

Having validated that PARprolink identifies PAR-binding proteins, we systematically investigated how endogenous proteins bind to different lengths of PAR. Emergent data suggest that signaling pathways are only activated when PAR length exceeds a certain threshold. Parthanatos, a PAR-dependent cell death pathway, is induced more strongly by long PAR (~60-mer) than short PAR (~15-mer).^33^ In addition, three DNA repair-related proteins (XPA, DEK, p53) preferentially bind long PAR, and the Chk1 kinase is only activated by long PAR.^34–36^ HeLa nuclear extract was cross-linked to either 8-mer or ~40-mer PAR photoaffinity probes. Comparing the intensities between these pull-downs uncovered 156 proteins that prefer ~40-mer PAR (Log_2_fold change > 2; Figure 3a and S5a; Supplementary Datafile). Importantly, we observed the long PAR-binding preference of DEK,^34^ validating that our approach identifies length-selective PAR binders.

**Figure 3.**
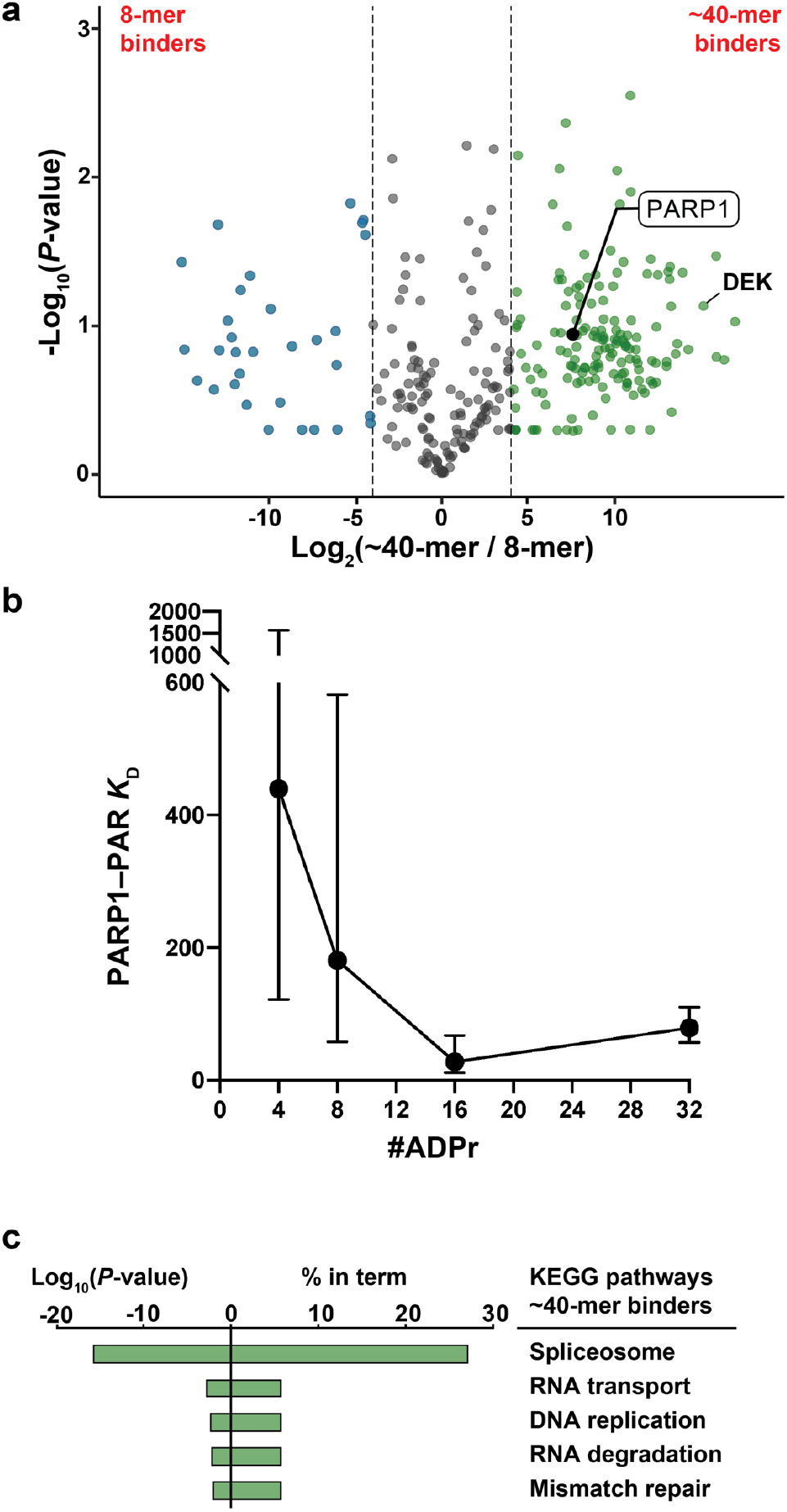
Defined-length probes reveal length-specific PAR-protein interactions. (a) Volcano plot of protein enrichment ratios and -Log_10_ P-values from proteomics experiments using either 8-or ~40-mer probes (n = 2). (b) The effect of PAR length on the affinity towards PARP1 measured with EMSA (the mean ± s.d.; *n* = 3). (c) Gene ontology analysis of proteins that were more abundant in the ~40-mer pull-down (enrichment ratio > 4).

Intriguingly, our analyses revealed that the central DNA repair protein PARP1 preferred binding to long PAR (~40-mer/8-mer = 13, Figure 3a). To verify this finding, EMSAs were performed with recombinant PARP1 and PAR of varied lengths (Figure 3b and S5b-c). We observed a 16-fold increase in PARP1-PAR affinity as PAR length increased from 4-to 16-mer. Importantly, the affinity of the PARP1-PAR interactions for the 16- and 32-mers (*K*_D_ = 11-110 nM) is in the same range as the reported affinities between PARP1 and nucleosomes (*K*_D_ = 2-100 nM).^37^ Given that PAR length is controlled temporally during DNA damage, where long polymers (>22-mer) are rapidly synthesized by PARP1 and then slowly degraded to shorter lengths,^27,38^ our data suggest that PAR length may control the dissociation of PARP1 from the chromatin during DNA repair.

Our investigation represents the first census of PAR binders. We took this opportunity to analyze global properties of the PAR-binding proteome. Gene ontology (GO) analysis on all 743 direct PAR binders (Figure 2) revealed the expected enrichment of several DNA repair pathways (Figure S6).^17^ Yet, the enrichments of physiological processes such as RNA splicing, RNA transport and DNA replication were even more significant. Notably, long PAR binders are enriched with proteins involved in nucleic acid metabolism, such as DNA repair and RNA splicing (Figure 3c). STRING protein association network analyses mapped two central cores—one involved in DNA repair and chromatin remodeling, and the other in RNA splicing and translation—along with distal clusters involved in metabolism and tRNA synthesis (Figure 4a and S7). Consistent with recent proteomic studies identifying ADP-ribosylated substrates,^10–16^ our analysis of the PAR-binding proteome strengthens the view that PAR has roles beyond DNA repair in metabolism and RNA regulation.^39,40^

**Figure 4.**
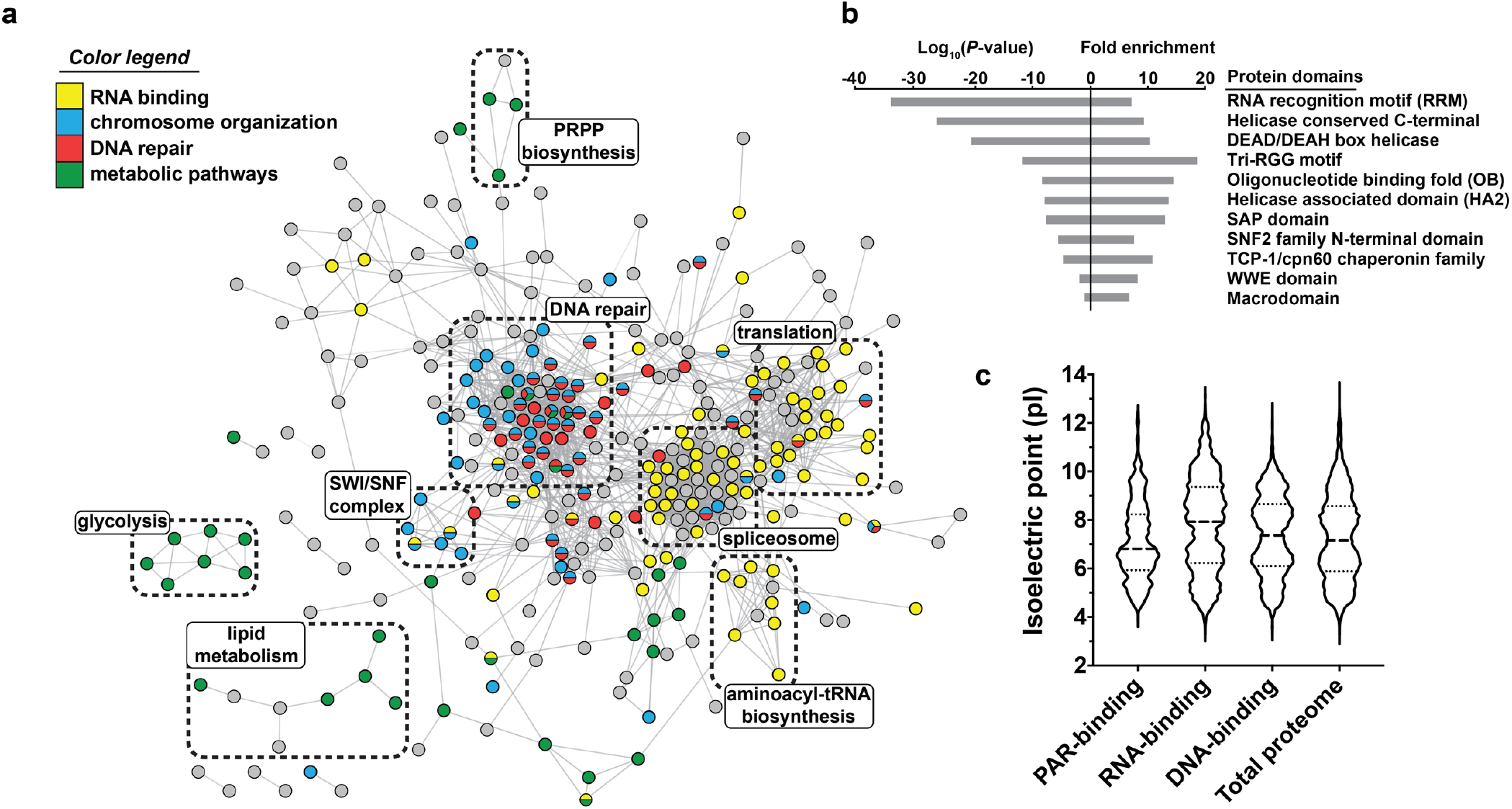
Global analyses of the human PAR-binding proteome. (a) STRING network analysis of high-confidence PAR-binding candidates (enrichment ratio > 8, *P* < 0.05, 416 genes) displaying at least one connection to another candidate. (b) Enrichment of known PAR-binding and other domains. (c) pI distribution among PAR-, RNA- and DNA-binding proteins.

We next assessed the enrichment of protein domains from the Pfam database in our dataset (Figure 4b and S8a-b; Supplementary Datafile).^41–43^ We observed a significant enrichment of multiple helicase-associated domains among direct PAR binders. Consistently, helicase activity is the most enriched molecular function based on GO analyses (Figure S6). Several known PAR-binding domains, such as WWE domain and macrodomain were also enriched. Amongst them, the most significant was the Tri-RGG motif, with 12 out of 16 Tri-RGG-containing proteins in the human proteome identified. In addition, we observed the enrichment of DNA/RNA-binding domains known to bind PAR, e.g., the RNA recognition motif and OB-fold.^44^ Therefore, it is not surprising that a significant amount of PAR binders are also known DNA-or RNA-binding proteins (*P* = 3.57 x 10^-6^, 2.00 x 10^-136^; Supplementary datafile).^21,22^ RNA- and DNA-binding proteins tend to have higher isoelectric points (median pI = 7.93 and 7.38). Unexpectedly, the median isoelectric point of PAR-binding proteins was lower than the proteome (pI = 6.81 vs 7.15, Figure 4c and S8c-f). Together, these data suggest a specific interaction between PAR and particular nucleic acid-binding domains, rather than a non-specific enrichment of positively-charged proteins.

In addition to defined motifs or domains, PAR-binding proteins were statistically enriched with proteins containing low-complexity sequence (*P* ≤ 5.5 x 10^-18^),^45,46^ which is critical for the formation of biomolecular condensates.^47^ Indeed, PAR-prolink identified PAR-binding proteins are enriched with components of biomolecular condensates such as DNA repair foci, nucleoli and stress granules (*P* = 8.36 x 10^-21^, 2.34 x 10^-57^, 5.97 x 10^-94^; Supplementary Datafile).^48–50^ Notably, PARy-lation of the DNA repair factor p53 and the nucleolar helicase DDX21 is dependent on their ability to bind PAR.^51,52^ Consistent with these studies, comparison with proteomics analyses of ADP-ribosylated substrates revealed that most direct PAR binders can also be ADP-ribosylated (647/743, 87%, *P* = 1.43 x 10^-246^; Supplementary datafile).^53^ Taken together, our data suggest that one or more PARylation events may trigger a wave of PAR binding-dependent PARylation in their vicinity, building extensive PAR-protein interaction networks to form biomolecular condensates in cells.^6,54^

This work describes the first proteomics method developed to identify direct PAR binders. Our census provides a global view of PARylation in DNA repair, RNA regulation and bio-molecular condensate formation, thereby serving as a rich resource to explore these frontiers in PAR biology.

## ASSOCIATED CONTENT

### Supporting Information

The Supporting Information is available free of charge on the ACS Publications website.

Experimental Methods, Figures S1-S8, NMR spectra of new compounds (PDF)

Proteomics Data and Analysis (XLSX)

## Notes

The authors declare no competing financial interest.

## ACKNOWLEDGMENT

We thank Dr. Srinivasan Yegnasubramanian for the UHRF1 plasmid, Dr. J. Paul Taylor for G3BP1 protein and plasmids for G3BP2, DDX6 and CAPRIN1, Dr. John Pascal for the PARP1 plasmid, and Tim Mitchison for in vitro synthesized PAR mixture. We thank the Johns Hopkins Discovery Award as well as NIH grants T32GM080189, T32GM008403, GM104135 (MMG), RF1AG071326 (AKLL) for funding.

## ABBREVIATIONS

RNF146: ring finger protein 146
BSA: bovine serum albumin
AK2: adenylate kinase 2
OB: oligonucleotide/oligosaccharide-binding

## REFERENCES

(1) Gibson, B.A.; Kraus, W. L. New Insights into the Molecular and Cellular Functions of poly(ADP-Ribose) and PARPs. Nat. Rev. Mol. Cell Biol. 2012, 13 (7), 411–424.

(2) Lüscher, B.; Bütepage, M.; Eckei, L.; Krieg, S.; Verheugd, P.; Shilton, B. H. ADP-Ribosylation, a Multifaceted Posttranslational Modification Involved in the Control of Cell Physiology in Health and Disease. Chem. Rev. 2018, 118 (3), 1092–1136.

(3) Suskiewicz, M. J.; Palazzo, L.; Hughes, R.; Ahel, I. Progress and Outlook in Studying the Substrate Specificities of PARPs and Related Enzymes. FEBS J. 2020. https://doi.org/10.1111/febs.15518.

(4) Sanderson, D. J.; Cohen, M. S. Mechanisms Governing PARP Expression, Localization, and Activity in Cells. Crit. Rev. Biochem. Mol. Biol. 2020, 1–14.

(5) Hottiger, M. O. SnapShot: ADP-Ribosylation Signaling. Mol. Cell 2015, 58 (6), 1134–1134.e1.

(6) Leung, A. K. L. Poly(ADP-Ribose): A Dynamic Trigger for Biomolecular Condensate Formation. Trends Cell Biol. 2020, 30 (5), 370–383.

(7) Lord, C. J.; Ashworth, A. PARP Inhibitors: Synthetic Lethality in the Clinic. Science 2017, 355 (6330), 1152–1158.

(8) Berger, N. A.; Besson, V. C.; Boulares, A. H.; Bürkle, A.; Chiarugi, A.; Clark, R. S.; Curtin, N. J.; Cuzzocrea, S.; Dawson, T. M.; Dawson, V. L.; Haskó, G.; Liaudet, L.; Moroni, F.; Pacher, P.; Radermacher, P.; Salzman, A. L.; Snyder, S. H.; Soriano, F. G.; Strosznajder, R. P.; Sümegi, B.; Swanson, R. A.; Szabó, C. Opportunities for the Repurposing of PARP Inhibitors for the Therapy of Non-Oncological Diseases. Br. J. Pharmacol. 2017, 175 (2), 192–222.

(9) Eustermann, S.; Wu, W.-F.; Langelier, M.-F.; Yang, J.-C.; Easton, L. E.; Riccio, A. A.; Pascal, J. M.; Neuhaus, D. Structural Basis of Detection and Signaling of DNA Single-Strand Breaks by Human PARP-1.Mol. Cell 2015, 60 (5), 742–754.

(10) Zhen, Y.; Yu, Y. Proteomic Analysis of the Downstream Signaling Network of PARP1. Biochemistry 2018, 57 (4), 429–440.

(11) Daniels, C. M.; Ong, S.-E.; Leung, A. K. L. The Promise of Proteomics for the Study of ADP-Ribosylation. Mol. Cell 2015, 58 (6), 911–924.

(12) Buch-Larsen, S. C.; Hendriks, I. A.; Lodge, J. M.; Rykær, M.; Furtwängler, B.; Shishkova, E.; Westphall, M. S.; Coon, J. J.; Nielsen, M. L. Mapping Physiological ADP-Ribosylation Using Activated Ion Electron Transfer Dissociation. Cell Rep. 2020, 32 (12), 108176.

(13) Nowak, K.; Rosenthal, F.; Karlberg, T.; Bütepage, M.; Thorsell, A.-G.; Dreier, B.; Grossmann, J.; Sobek, J.; Imhof, R.; Lüscher, B.; Schüler, H.; Plückthun, A.; Leslie Pedrioli, D. M.; Hottiger, M. O. Engineering Af1521 Improves ADP-Ribose Binding and Identification of ADP-Ribosylated Proteins. Nat. Commun. 2020, 11 (1), 5199.

(14) Carter-O’Connell, I.; Jin, H.; Morgan, R. K.; David, L. L.; Cohen, M. S. Engineering the Substrate Specificity of ADP-ribosyltransferases for Identifying Direct Protein Targets. J. Am. Chem. Soc. 2014, 136 (14), 5201–5204.

(15) Jiang, H.; Kim, J. H.; Frizzell, K. M.; Kraus, W. L.; Lin, H. Clickable NAD Analogues for Labeling Substrate Proteins of Poly(ADP-ribose) Polymerases. J. Am. Chem. Soc. 2010, 132 (27), 9363–9372.

(16) Gibson, B. A.; Zhang, Y.; Jiang. H.; Hussey, K.; Shrimp, J. H.; Lin, H.; Schwede, F.; Yu, Y.; Kraus, W. L Chemical genetic discovery of PARP targets reveals a role for PARP-1 in transcription elongation. Science 2016, 6294 (353), 45–50.

(17) Gagné, J.-P.; Pic, E.; Isabelle, M.; Krietsch, J.; Ethier, C.; Paquet, E.; Kelly, I.; Boutin, M.; Moon, K.-M.; Foster, L. J.; Poirier, G. G. Quantitative Proteomics Profiling of the poly(ADP-Ribose)-Related Response to Genotoxic Stress. Nucleic Acids Res. 2012, 40 (16), 7788–7805.

(18) Wright, R. H. G.; Lioutas, A.; Le Dily, F.; Soronellas, D.; Pohl, A.; Bonet, J.; Nacht, A. S.; Samino, S.; Font-Mateu, J.; Vicent, G. P.; Wierer, M.; Trabado, M. A.; Schelhorn, C.; Carolis, C.; Macias, M. J.; Yanes, O.; Oliva, B.; Beato, M. ADP-Ribose-Derived Nuclear ATP Synthesis by NUDIX5 Is Required for Chromatin Remodeling. Science 2016, 352 (6290), 1221–1225.

(19) Teloni, F.; Altmeyer, M. Readers of poly(ADP-Ribose): Designed to Be Fit for Purpose. Nucleic Acids Res. 2016, 44 (3), 993–1006.

(20) Krietsch, J.; Rouleau, M.; Pic, É.; Ethier, C.; Dawson, T. M.; Dawson, V. L.; Masson, J.-Y.; Poirier, G. G.; Gagné, J.-P. Reprogramming Cellular Events by poly(ADP-Ribose)-Binding Proteins. Mol. Aspects Med. 2013, 34 (6), 1066–1087.

(21) Gerstberger, S.; Hafner, M.; Tuschl, T. A Census of Human RNA-Binding Proteins. Nat. Rev. Genet. 2014, 15 (12), 829–845.

(22) Lambert, S. A.; Jolma, A.; Campitelli, L. F.; Das, P. K.; Yin, Y.; Albu, M.; Chen, X.; Taipale, J.; Hughes, T. R.; Weirauch, M. T. The Human Transcription Factors. Cell 2018, 175 (2), 598–599.

(23) Castello, A.; Fischer, B.; Eichelbaum, K.; Horos, R.; Beckmann, B. M.; Strein, C.; Davey, N. E.; Humphreys, D. T.; Preiss, T.; Steinmetz, L. M.; Krijgsveld, J.; Hentze, M. W. Insights into RNA Biology from an Atlas of Mammalian mRNA-Binding Proteins. Cell 2012, 149 (6), 1393–1406.

(24) Arguello, A. E.; DeLiberto, A. N.; Kleiner, R. E. RNA Chemical Proteomics Reveals the N6-Methyladenosine (m6A)-Regulated Protein-RNA Interactome. J. Am. Chem. Soc. 2017, 139 (48), 17249–17252.

(25) Tan, E. S.; Krukenberg, K. A.; Mitchison, T. J. Large Scale Preparation and Characterization of poly(ADP-Ribose) and Defined Length Polymers. Anal. Biochem. 2012.

(26) Ando, Y.; Elkayam, E.; McPherson, R. L.; Dasovich, M.; Cheng, S.-J.; Voorneveld, J.; Filippov, D. V.; Ong, S.-E.; Joshua-Tor, L.; Leung, A. K. L. ELTA: Enzymatic Labeling of Terminal ADP-Ribose. Mol. Cell 2019, 73 (4), 845–856.e5.

(27) Spitale, R. C.; Crisalli, P.; Flynn, R. A.; Torre, E. A.; Kool, E. T.; Chang, H. Y. RNA SHAPE Analysis in Living Cells. Nat. Chem. Biol. 2013, 9 (1), 18–20.

(28) Mortimer, S. A.; Weeks, K. M. Time-resolved RNA SHAPE Chemistry. J. Am. Chem. Soc. 2008, 130 (48), 16178–16180.

(29) Dormán, G.; Prestwich, G. D. Benzophenone photophores in Biochemistry. Biochemistry 1994, 33 (19), 5661–5673.

(30) Wong, I.; Lohman, T. M. A Double-Filter Method for Nitro-cellulose-Filter Binding: Application to Protein-Nucleic Acid Interactions. Proc. Natl. Acad. Sci. U. S. A. 1993, 90 (12), 5428–5432.

(31) Cox, J.; Hein, M. Y.; Luber, C. A.; Paron, I.; Nagaraj, N.; Mann, M. Accurate Proteome-Wide Label-Free Quantification by Delayed Normalization and Maximal Peptide Ratio Extraction, Termed MaxLFQ. Mol. Cell. Proteomics 2014, 13 (9), 2513–2526.

(32) Li, M.; Lu, L.-Y.; Yang, C.-Y.; Wang, S.; Yu, X. The FHA and BRCT Domains Recognize ADP-Ribosylation during DNA Damage Response. Genes Dev. 2013, 27 (16), 1752–1768.

(33) Andrabi, S. A.; Kim, N. S.; Yu, S.-W.; Wang, H.; Koh, D. W.; Sasaki, M.; Klaus, J. A.; Otsuka, T.; Zhang, Z.; Koehler, R. C.; Hurn, P. D.; Poirier, G. G.; Dawson, V. L.; Dawson, T. M. Poly(ADP-Ribose) (PAR) Polymer Is a Death Signal. Proc. Natl. Acad. Sci. U. S. A. 2006, 103 (48), 18308–18313.

(34) Fahrer, J.; Kranaster, R.; Altmeyer, M.; Marx, A.; Bürkle, A. Quantitative Analysis of the Binding Affinity of poly(ADP-Ribose) to Specific Binding Proteins as a Function of Chain Length. Nucleic Acids Res. 2007, 35 (21), e143.

(35) Fahrer, J.; Popp, O.; Malanga, M.; Beneke, S.; Markovitz, D. M.; Ferrando-May, E.; Bürkle, A.; Kappes, F. High-Affinity Interaction of poly(ADP-Ribose) and the Human DEK Oncoprotein Depends upon Chain Length. Biochemistry 2010, 49 (33), 7119–7130.

(36) Min, A.; Im, S.-A.; Yoon, Y.-K.; Song, S.-H.; Nam, H.-J.; Hur, H.-S.; Kim, H.-P.; Lee, K.-H.; Han, S.-W.; Oh, D.-Y.; Kim, T.-Y.; O’Connor, M. J.; Kim, W.-H.; Bang, Y.-J. RAD51C-Deficient Cancer Cells Are Highly Sensitive to the PARP Inhibitor Olaparib. Mol. Cancer Ther. 2013, 12 (6), 865–877.

(37) Wielckens, K.; Schmidt, A.; George, E.; Bredehorst, R.; Hilz, H. DNA Fragmentation and NAD Depletion. Their Relation to the Turnover of Endogenous mono(ADP-Ribosyl) and poly(ADP-Ribosyl) Proteins. J. Biol. Chem. 1982, 257 (21), 12872–12877.

(38) Muthurajan, U. M.; Hepler, M. R. D.; Hieb, A. R.; Clark, N. J.; Kramer, M.; Yao, T.; Luger, K. Automodification Switches PARP-1 Function from Chromatin Architectural Protein to Histone Chaperone. Proc. Natl. Acad. Sci. U. S. A. 2014.

(39) Kim, D.-S.; Challa, S.; Jones, A.; Kraus, W. L. PARPs and ADP-Ribosylation in RNA Biology: From RNA Expression and Processing to Protein Translation and Proteostasis. Genes Dev. 2020, 34 (5-6), 302–320.

(40) Szántó, M.; Bai, P. The Role of ADP-Ribose Metabolism in Metabolic Regulation, Adipose Tissue Differentiation, and Metabolism. Genes Dev. 2020, 34 (5-6), 321–340.

(41) Altmeyer, M.; Neelsen, K. J.; Teloni, F.; Pozdnyakova, I.; Pellegrino, S.; Grøfte, M.; Rask, M.-B. D.; Streicher, W.; Jungmichel, S.; Nielsen, M. L.; Lukas, J. Liquid Demixing of Intrinsically Disordered Proteins Is Seeded by poly(ADP-Ribose). Nat. Commun. 2015, 6, 8088.

(42) Gupte, R.; Liu, Z.; Kraus, W. L. PARPs and ADP-Ribosylation: Recent Advances Linking Molecular Functions to Biological Outcomes. Genes Dev. 2017, 31 (2), 101–126.

(43) Finn, R. D.; Tate, J.; Mistry, J.; Coggill, P. C.; Sammut, S. J.; Hotz, H.-R.; Ceric, G.; Forslund, K.; Eddy, S. R.; Sonnhammer, E. L. L.; Bateman, A. The Pfam Protein Families Database. Nucleic Acids Res. 2008, 36 (Database issue), D281–D288.

(44) Zhang, F.; Chen, Y.; Li, M.; Yu, X. The Oligonucleotide/oli-gosaccharide-Binding Fold Motif Is a poly(ADP-Ribose)-Binding Domain That Mediates DNA Damage Response. Proc. Natl. Acad. Sci. U. S. A. 2014, 111 (20), 7278–7283.

(45) March, Z. M.; King, O. D.; Shorter, J. Prion-like Domains as Epigenetic Regulators, Scaffolds for Subcellular Organization, and Drivers of Neurodegenerative Disease. Brain Res. 2016, 1647, 9–18.

(46) Han, T. W.; Kato, M.; Xie, S.; Wu, L. C.; Mirzaei, H.; Pei, J.; Chen, M.; Xie, Y.; Allen, J.; Xiao, G.; McKnight, S. L. Cell-free Formation of RNA Granules: Bound RNAs Identifiy Features and Components of Cellular Assemblies. Cell 2012, 149 (4), 768–779.

(47) Banani, S. F.; Lee, H. O.; Hyman, A. A.; Rosen, M. K. Bio-molecular Condensates: Organizers of Cellular Biochemistry. Nat. Rev. Mol. Cell Biol. 2017, 18 (5), 285–298.

(48) Chou, D. M.; Adamson, B.; Dephoure, N. E.; Tan, X.; Nottke, A. C.; Hurov, K. E.; Gygi, S. P.; Colaiácovo, M. P.; Elledge, S. J. A Chromatin Localization Screen Reveals Poly(ADP-ribose)-regulated Recruitment Polycomb and NuRD Complexes to Sites of DNA Damage. Proc. Natl. Acad. Sci. U. S. A. 2010, 107 (43), 18475–18480.

(49) Ahmad, Y.; Boisvert, F.; Gregor, P.; Cobley, A.; Lamond, A. I. NOPdb: Nucleolar Proteome Database—2008 Update. Nucleic Acids Res. 2009, 37 (D1), D181–D184.

(50) Nunes, C.; Mestre, I.; Marcelo, A.; Koppenol, R.; Matos. C. A.; Nóbrega, C.. MSGP: the first database of the protein components of the mammalian stress granules. Database. 2019, 1–7.

(51) Fischbach, A.; Krüger, A.; Hampp, S.; Assmann, G.; Rank, L.; Hufnagel, M.; Stöckl, M. T.; Fischer, J. M. F.; Veith, S.; Rossatti, P.; Ganz, M.; Ferrando-May, E.; Hartwig, A.; Hauser, K.; Wiesmüller, L.; Bürkle, A.; Mangerich, A. The C-Terminal Domain of p53 Orchestrates the Interplay between Non-Covalent and Covalent poly(ADP-Ribosyl)ation of p53 by PARP1. Nucleic Acids Res. 2018, 46 (2), 804–822.

(52) Kim, D.-S.; Camacho, C. V.; Nagari, A.; Malladi, V. S.; Challa, S.; Kraus, W. L. Activation of PARP-1 by snoRNAs Controls Ribosome Biogenesis and Cell Growth via the RNA Helicase DDX21.Mol. Cell 2019, 75 (6), 1270–1285.e14.

(53) Ayyappan, V.; Wat, R.; Barber, C.; Vivelo, C. A.; Gauch, K.; Visanpattanasin, P.; Cook, G.; Sazeides, C.; Leung, A. K. L. ADPriboDB 2.0: an updated database of ADP-ribosylated proteins. Nucleic Acids Res. 2020, 1, 1–5.

(54) Dasovich, M.; Leung, A. K. L. A Nucleolar PARtnership Expands PARP Roles in RNA Biology and the Clinical Potential of PARP Inhibitors. Mol. Cell 2019, 75 (6), 1089–1091.

