## Supporting information for "PARprolink: a photoaffinity probe for identifying poly(ADP-ribose)-binding proteins"

### PARprolink: a photoaffinity probe to identify direct poly(ADP-ribose) binders

Table of Contents

S2-16. Methods
S17. Figure S1
S18. Figure S2
S19. Figure S3
S20. Figure S4
S21. Figure S5
S22. Figure S6
S23. Figure S7
S24. Figure S8
S25-28. NMR spectra of new compounds

#### General methods

Solvents were distilled prior to use as follows: tetrahydrofuran (THF) was distilled from sodium benzophenone ketyl; pyridine and dichloromethane were distilled from calcium hydride. All commercial reagents were used without further purification unless otherwise stated. All glassware was dried at 120 °C for 24 hr prior to use. All reactions were carried out under a positive pressure of argon atmosphere and monitored by TLC on silica gel G-25 UV254 (0.25 mm). Spots were detected under UV light. Column flash chromatography was performed with Silicycle grade 70–230 mesh, 60–200 µm, 60 Å silica. The ratio between silica gel and crude product ranged from 100:1 to 60:1 (w/w).

$^{1}H$ (400 MHz) and $^{13}C$ (101 MHz) NMR spectra were recorded on a 400 MHz NMR spectrometer. All spectra were recorded in deuterated chloroform ($CDCl_3$), deuterated methanol ($CD_3OD$) or deuterated DMSO ($d_6$-DMSO). Chemical shifts ($\delta_H$ and $\delta_C$) are reported in parts per million (ppm) and coupling constants are expressed in hertz (Hz). Splitting patterns in 1H NMR spectra are designated as s (singlet), dd (doublet of doublets), and m (multiplet).

#### Methyl 2-chloromethylnicotinate

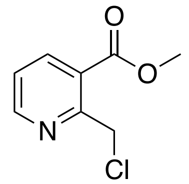


The compound was prepared according to literature procedures.[1] Briefly, a round-bottom flask equipped with stirring was flushed with argon. Methyl 2-methylnicotinate (1.6 g, 10.6 mmol) was added via syringe, followed by dichloromethane (8 mL). Trichloroisocyanuric acid (3.7 g, 15.9 mmol) was added and the mixture was stirred at ambient temperature overnight. The reaction was diluted with dichloromethane (20 mL), then saturated sodium bicarbonate (10 mL) was added slowly. The aqueous and organic layers were separated. The organic layer was washed twice with brine (2 x 10 mL), then dried over sodium sulfate, filtered and concentrated to afford the title compound as a yellow oil (1.687 g, 86%). Observed chemical shifts agree with literature reports.

**2-Chloromethylnicotinic acid**

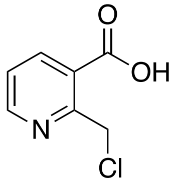

Methyl 2-chloromethylnicotinate (0.5 g, 2.7 mmol) was stirred in 5 mL of 1:1 MeOH:10% aq. NaOH at ambient temperature in a round-bottom flask equipped with stirring. After 1 hr, TLC indicated complete consumption of starting material. The crude reaction was washed once with ether (5 mL), then acidified to pH 4 with 10% aq. HCl and extracted with ethyl acetate (5 x 10 mL). The ethyl acetate dried over sodium sulfate, filtered and concentrated to afford the title compound as a light brown powder: $^1$H NMR (400 MHz, $CD_3OD$) $\delta$ 8.80 (dd, $J$ = 1.8, 4.8, 1H), 8.40 (dd, $J$ = 1.8, 7.9, 1H), 7.42 (dd, $J$ = 4.9, 7.9, 1H), 5.17 (s, 1H); $^{13}$C {$^1$H} NMR (101 MHz, $CD_3OD$) $\delta$ 166.8, 156.2, 151.9, 139.0, 126.2, 123.8; HRMS (ESI–TOF) $m/z$ (M + H)$^+$ calcd for $C_7H_6ClNO_2$ = 172.016532, found $m/z$ = 172.0165

**(4-(13-Hydroxy-2,5,8,11-tetraoxatridecyl)phenyl)(phenyl)methanone**

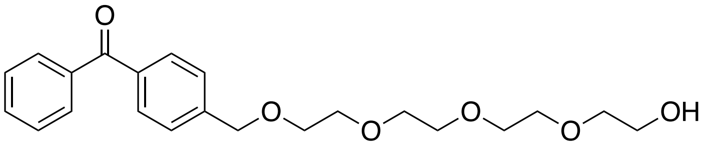

A round-bottom flask equipped with stirring was flushed with argon. Tetraethylene glycol (3.88 g, 20 mmol) was added via syringe was azeotropically dried with anhydrous pyridine (2 $\times$ 4.0 mL), redissolved in anhydrous THF (10 mL) and cooled to 0 °C. Sodium hydride (0.176 g, 4.4 mmol) was added. The mixture was stirred at 0 °C for 30 min, then 4-(bromomethyl) benzophenone (1.1 g, 4 mmol) was added. The mixture was stirred at 0 °C for 30 min, then removed from ice and stirred overnight at ambient temperature. Saturated aqueous ammonium chloride (5 mL) was added slowly to quench the reaction. The organic solvent was removed under reduced pressure. The slurry was diluted with brine (10 mL) and then extracted with dichloromethane (5 $\times$ 10 mL). The organic layer dried over sodium sulfate, filtered and concentrated under reduced pressure. The crude product was purified by chromatography (0-10% methanol in 1:1 ethyl acetate:dichloromethane), yielding the title compound as a light brown oil (1.0 g, 32.6 mmol, 65%): $^1$H NMR (400 MHz, $CDCl_3$) $\delta$ 7.82–7.75 (m, 4H), 7.62–7.56 (m, 1H), 7.51–7.44 (m, 4H), 4.65 (s, 2H), 3.76–3.6 (m, 16H); $^{13}$C {$^1$H} NMR (101 MHz, $CDCl_3$) $\delta$ 195.8, 142.9, 137.2, 136.2, 131.9, 129.7, 129.5, 127.8, 126.7, 77.3, 77.0, 76.6, 72.1, 70.1, 70.1, 70.1, 69.8, 69.4, 61.1; HRMS (EI) *m/z* $(M)^+$ calcd for $C_{22}H_{28}O_6$ = 388.188590, found *m/z* = 388.18828

**2-(15-(4-Benzoylphenyl)-2,5,8,11,14-pentaoxapentadecyl)nicotinic acid**

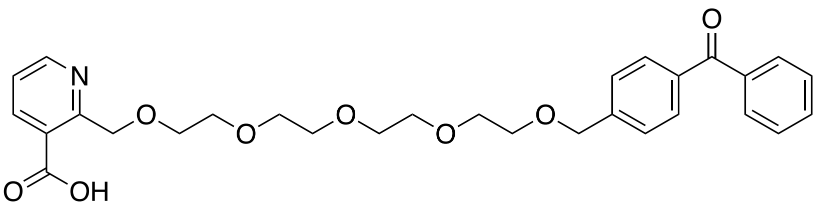


A round-bottom flask equipped with stirring was flushed with argon. (4-(13-hydroxy-2,5,8,11-tetraoxatridecyl)phenyl)(phenyl)methanone (0.388 g, 1 mmol) was added via syringe, azeotropically dried with pyridine (2 $\times$ 4 mL), redissolved in anhydrous THF (10 mL) and cooled to 0 °C. Sodium hydride (0.13 g, 3.3 mmol) was added. The mixture was stirred at 0 °C for 30 min, then 2-(chloromethyl) nicotinate (0.26 g, 2.5 mmol) was added. The mixture was stirred at 0 °C for 30 min, then removed from ice and stirred overnight at ambient temperature.

Saturated aqueous sodium bicarbonate (5 mL) was added slowly to quench the reaction. The organic solvent was removed under reduced pressure. The aqueous slurry was washed with diethyl ether (10 mL) and then acidified to pH 4 with concentrated HCl. The aqueous layer was extracted with ethyl acetate (5 × 10 mL). The combined ethyl acetate layers were dried over sodium sulfate, filtered and concentrated under reduced pressure. The crude product was purified by column chromatography using 5-20% methanol in dichloromethane, yielding the title compound as a brown oil (0.29 g, 0.55 mmol, 55%) $^1$H NMR (400 MHz, $CD_3OD$) $\delta$ 8.59 (1H, dd, $J$ = 1.7, 4.9), 8.23 (dd, 1H, $J$ = 1.7, 7.8), 7.75 (m, 4H), 7.62 (m, 1H), 7.52 (m, 5H), 7.44 (dd, 1H, $J$ = 5, 7.8), 4.99 (s, 2H), 4.64 (s, 2H), 3.65 (m, 16H); $^{13}$C {$^1$H} NMR (101 MHz, $CD_3OD$) $\delta$ 198.3, 158.4, 150.8, 145.1, 140.5, 139.0, 138.0, 133.9, 131.3, 131.1, 129.6, 128.5, 124.4, 73.4, 73.2, 71.6, 71.5, 71.4, 71.4, 71.3, 71.0, 50.0, 49.7, 49.6, 49.4, 49.1, 48.9, 48.5; HRMS (ESI–TOF) $m/z$ (M + H)$^+$ calcd for $C_{29}H_{33}NO_8$ = 524.228444 found $m/z$ = 524.2290

**(4-(15-(3-(1*H*-imidazole-1-carbonyl)pyridin-2-yl)-2,5,8,11,14-pentaoxapentadecyl)phenyl)(phenyl)methanone**

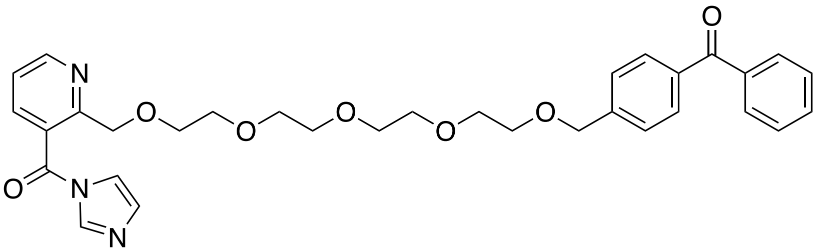

Anhydrous $d_6$-DMSO was prepared by drying over 4 Å sieves. 2-(15-(4-Benzoylphenyl)-2,5,8,11,14-pentaoxapentadecyl)nicotinic acid (60 μL, 1 M in anhydrous $d_6$-DMSO) and carbonyldiimidazole (60 μL, 2 M in anhydrous $d_6$-DMSO) were combined in a 0.5 mL conical vial equipped with stirring. To monitor reaction progress with $^1$H NMR, a small aliquot was taken and diluted further in $d_6$-DMSO. Product formation was measured by integrating the shift of a methylene peak (see $^1$H NMR spectrum on page S28 for an example).The mixture was stirred for 1 hr at ambient temperature, sealed with parafilm, then stored at -80 °C. This compound was not isolated. The crude product was used as a 0.5 M stock in the reaction with PAR. The stock should always be under argon atmosphere and warmed up to ambient temperature before use.

**Labeling poly(ADP-ribose) with dATP-biotin using OAS1**

PAR was synthesized and HPLC-fractionated to defined length according to literature precedent.[2,3] PAR of defined length (3–20 nmol depending on the reaction, ≤100 μM) was reacted with biotin-dATP (Jena BioScience, 3-fold molar excess to PAR), OAS1 (20:1 molar ratio of PAR:OAS1) and poly(I:C)(low MW)(1:1 ratio with OAS1, w/w) in 20 mM Tris pH 7.5, 20 mM $Mg(OAc)_2$, 2.5 mM DTT in a 125 μL or 250 μL reaction, depending on the scale. The reaction was incubated at 37 °C / 750 rpm for 2 hr then purified with DHBB-agarose.

**Purification of PAR with DHBB-agarose**

*m*-Aminophenylboronic acid–agarose (Sigma, 100 μL of suspension for 20 nmol PAR, 1 BV) was equilibrated with 10 bead volume (BV) of AAEG9 buffer (0.25 M ammonium acetate pH 9.0, 6 M guanidine-HCl, 10 mM EDTA) for 1 min with end-over end rotation. The beads were pelleted with 500 *g* for 2 min and the supernatant was removed. The beads were equilibrated with 10 BV of $ddH_2O$ for 1 min with end-over end rotation. The beads were pelleted with 500 *g* for 2 min and the supernatant was removed. The beads were equilibrated with 10 BV of AAEG9 buffer (0.25 M ammonium acetate pH 9.0, 6 M guanidine-HCl, 10 mM EDTA) for 1 min with end-over end rotation. The beads were pelleted with 500 *g* for 2 min and the supernatant was removed. The reaction containing PAR was diluted with 9 volumes of AAEG9 buffer, then added to the beads and incubated for 1 hr at R.T. with end-over-end rotation. The bead-PAR mixture was transferred to a 10 mL disposable gravity column (Bio-

Rad). The beads were washed with 10 BV AAEG9 buffer, followed by 10 BV 1 M ammonium acetate pH 9.0. The PAR was eluted with 37 °C dd$H_2O$ (10 x 1 BV). Eluates with absorbance at 260 nm were combined, flash frozen, lyophilized, then resuspended in dd$H_2O$. A UV-vis spectrum was collected to estimate [PAR].

**Acylation of biotin-PAR with (4-(15-(3-(1*H*-imidazole-1-carbonyl)pyridin-2-yl)-2,5,8,11,14-pentaoxapentadecyl)phenyl)(phenyl)methanone**

Biotin-PAR (50-100 µM) was mixed with (4-(15-(3-(1*H*-imidazole-1-carbonyl)pyridin-2-yl)-2,5,8,11,14-pentaoxapentadecyl)phenyl)(phenyl)methanone (50 mM) in 0.1 M HEPES pH 8.0, 0.1 M NaCl, 6 mM $MgCl_2$ in a reaction with a total volume of 50 µL. The mixture was incubated at ambient temperature for 3 hr with constant vortexing. The pH of the reaction becomes acidic over time, so aqueous sodium hydroxide (1 µL of 1 M solution) was added periodically to maintain a pH of ~8. After completion, the PAR was purified with DHBB-agarose as described above.

**Purification of acylated PAR with HPLC**

An Agilent Infinity II HPLC was loaded with 0.1 M TEAA pH 7.5 and acetonitrile into the appropriate solvent ports. The detector was set to record 215 nm and 258 nm. An Infinity Poroshell 120 EC-C18 (Agilent # 693970-902T) was connected and equilibrated with 95% TEAA pH 7.5 and 5% acetonitrile. Aliquots of acylated biotin-PAR (up to 20 nmol) were injected and separated with a gradient (0-10 min, 5% acetonitrile; 10-40 min, 5%-40% acetonitrile). Fractions (0.25 mL) were automatically collected during the entire run. Strongly absorbing fractions were combined based on the number of benzophenones in 1.7 mL tubes. Combined fractions were flash frozen in liquid nitrogen, then lyophilized to dryness. The PAR was resuspended with 50 µL of TE buffer. A UV-vis spectrum was collected to estimate [PARprolink] with the equation given below.

##### Determining PARprolink concentration

PAR concentration was determined using the following equation: [PAR] = $[(A_{258})\ cm^{-1}]/([13{,}500\ cm^{-1}\ M^{-1}] \times n + [18{,}000\ cm^{-1}\ M^{-1}] \times m)$ where $n$ = # of adenines and $m$ = # of benzophenones.

##### Western blot

For each blot, Ponceau S (Sigma-Aldrich #P7170) was used according to the manufacturer's instructions. The following antibodies were used: IRDye 800 CW Streptavidin (Li-Cor #926-32230, 1:3,000 in TBS + 0.15% v/v Tween-20 + 0.06% SDS + 0.02% sodium sodum azide), anti-FLAG (Sigma-Aldrich #F7425, 1:1,000 in TBS-T + 0.02% sodium azide), IRDye 680LT (Li-Cor #926-68021, 1:20,000 in TBS-T).

##### Cross-linking with recombinant protein

All photolyses were carried out in 1.7 mL or 0.6 mL clear tubes in a Rayonet photoreactor fitted with 16 lamps with maximum output at 350 nm. Photolysis was performed at room temperature on a rotating platform. PARprolink (15-mer, 100 nM) was mixed with the indicated concentration of RNF146-WWE-FLAG or BSA in binding buffer (20 mM HEPES pH 7.5, 100 mM NaCl) in a reaction with a total volume of 30 µL. Reactions were equilibrated on ice for 30 min, then photolyzed for 10 min. Reactions were mixed with 4X SDS sample loading buffer, incubated at 95 °C for 5 min, then separated with SDS-PAGE. Proteins were transferred to nitrocellulose, blocked in 5% non-fat dry milk in TBS for 1 h, then blotted with

IRdye 800CW Streptavidin for 1 hr before the data were acquired with an Odyssey infrared imager (Li-Cor).

**Cell culture**

HeLa cells were cultured at 37 °C in a humidified atmosphere with 5% $CO_2$ in DMEM (Life Technologies) supplemented with 10% FBS (HyClone).

**Preparation of whole-cell extracts**

A 10 cm dish was seeded with 1 million HeLa cells in DMEM + 10% FBS and grown at 37 °C + 5% $CO_2$ for 24 hr prior to harvesting. Cells were washed with ice-cold PBS (3 x 10 mL), then 1 mL of ice-cold NP-40 lysis buffer (50 mM Tris pH 7.5, 150 mM NaCl, 5 mM $MgCl_2$, 1 mM PMSF, 1% v/v NP-40, 1X SIGMAFAST protease inhibitor cocktail) was added. The dish was incubated flat on ice for 15 min, the cells were then scraped into a pre-chilled 1.7 mL tube and incubated at 4 °C with end-over-end rotation for 15 min. The lysate was clarified by spinning at 20,000 *g* / 4 °C for 10 min. Protein concentration was determined with a Bradford assay (ThermoFisher) using NP-40 lysis buffer as the diluent for the BSA standards.

**Cross-linking with recombinant protein in whole-cell extract**

All photolyses were carried out in 1.7 mL or 0.6 mL clear tubes in a Rayonet photoreactor fitted with 16 lamps with maximum output at 350 nm. Photolysis was performed at room temperature on a rotating platform. PARprolink (15-mer, 100 nM) was mixed with 7.2 µM or the indicated concentration of RNF146-WWE-FLAG in lysis buffer (50 mM Tris pH 7.5, 150 mM NaCl, 5 mM $MgCl_2$, 1 mM PMSF, 1X SIGMAFAST protease inhibitor cocktail) in a reaction with a total volume of 40 µL. For the “+unlabeled PAR” reaction, 30 µM of 15-mer PAR was also added. Reactions were equilibrated on ice for 60 min, then photolyzed for 10 min. Reactions were mixed with 4X SDS sample loading buffer, incubated at 95 °C for 5 min, then separated with SDS-PAGE. Proteins were transferred to nitrocellulose, blocked in 5% non-fat dry milk in TBS for 1 hr, then blotted with IRdye 800CW Streptavidin for 1 hr, before the data were acquired with an Odyssey infrared imager (Li-Cor). The membrane was then incubated with anti-FLAG and anti-Rabbit 680LT, each at room temperature for 1 hr, before the data were acquired with an Odyssey infrared imager (Li-Cor).

**Preparation of HeLa nuclear extracts**

Extracts were prepared as described,[13] with modifications. HEPES pH 8.0 was used instead of Tris pH 7.9, 1 mM DTT was added to the low-salt and high-salt buffers, and 1X SIGMAFAST protease inhibitor cocktail was used in all buffers except for the dialysis buffer, where 0.2 mM PMSF was used.

**Cross-linking and pull-down in nuclear extract**

All photolyses were carried out in 1.7 mL clear tubes in a Rayonet photoreactor fitted with 16 lamps with maximum output at 350 nm. Photolysis was done on ice in a glass dewar. Care was taken to ensure all tubes equidistant from the UV lamps. Nuclear extract (3 mg/mL) in extract buffer (20 mM HEPES pH 8.0, 100 mM KCl, 0.2 mM EDTA, 1 mM DTT, 20% glycerol, 0.2 mM PMSF) supplemented with PARG inhibitor (PDD 00017273, 100 μM) was incubated on ice for 30 min before the addition of PARprolink (8-mer or ~40-mer, 2 μM) in a reaction with a total volume of 500 μL. Reactions were equilibrated on ice for 30 min, then photolyzed for 20 min. Reactions were diluted to 0.1 mg/mL with extract buffer in 50 mL tubes, then streptavidin sepharose (60 μL) was added and the mixtures were incubated with end-over-end rotation for 3 hr. The streptavidin sepharose was sedimented by spinning at 1,000 *g* and the supernatant was removed. The streptavidin sepharose was resuspended in 0.5 mL

TBS, then transferred to 1.7 mL tubes and washed with SDS wash buffer (5 x 1 mL, 100 mM Tris pH 8, 1% SDS, 250 mM NaCl), urea wash buffer (5 x 1 mL, 100 mM Tris pH 8, 8 M urea), 20% acetonitrile (5 x 1 mL) and TBS (2 x 1 mL), in that order. For each wash, the tubes were incubated with end-over-end rotation for 1 min at ambient temperature, then the Sepharose was pelleted with 500 *g* for 2 min and the supernatant was removed.

**On-bead digestion and peptide desalting**

The streptavidin sepharose was resuspended in denaturing buffer (100 µL, 50 mM Tris pH 7.8, 8 M urea) and incubated with tris,(2-carboxyuethyl)phosphine (TCEP, 1 mM) at 37 °C / 1,400 rpm for 20 min. 2-chloroacetamide (CAM , 2 mM) was added and samples were incubated) at 37 °C / 1,400 rpm for 20 min. TCEP was increased to 2 mM and the samples were incubated for an additional 20 min at 37 °C / 1,400 rpm. Urea was diluted to 4 M with 25 mM ammonium bicarbonate. LysC (1 µg) was added to each sample and the samples were incubated for 2 hr at 37 °C / 1,400 rpm. Urea was diluted to 2 M with 25 mM ammonium bicarbonate, then trypsin (1 µg) was added to each sample and the samples were incubated overnight at 37 °C / 1,400 rpm. Each step of desalting was performed by spinning at 500 g. StageTips were conditioned with methanol (50 µL) followed by StageTip buffer (5% acetonitrile, 0.1% trifluoroacetic acid). The streptavidin sepharose was pelleted by spinning at 1,000 *g* then the supernatant was applied to the StageTip. The StageTips were washed with StageTip buffer (50 µL) then submitted for LC-MS analysis.

**nanoLC-MS/MS analyses**

Peptide samples were separated on a Thermo Easy-nLC1200 UHPLC instrument (Odense, DK) using 20 cm long fused silica capillary columns (100 µm ID) packed with 3 µm 120 Å reversed phase C18 beads (Dr. Maisch, Ammerbuch, DE). The LC gradient was 90 min with 4−32% B at 300 nL/min. LC solvent A was 0.1% aq. acetic acid and LC solvent B was 0.1% acetic acid, 80% acetonitrile. MS data was collected with a Thermo Fisher Scientific Orbitrap Fusion Lumos Tribrid spectrometer using data-dependent analysis with a 2 sec cycle time for MS1 acquisition and HCD-MS2 collected in the Orbitrap by Top15 precursor selection.

**Mass Spectrometry data analysis**

Under “raw data” the .raw files were loaded into MaxQuant version 1.6.2.6. Each file was labeled as its own experiment. Under “group-specific parameters”, multiplicity was set to 1, enzymes were set to LysC and trypsin, the variable modifications acetyl (N-term) and oxidation (M) were allowed, and the carbamidomethylation (C) was the only fixed modification. Under “global parameters”, the FASTA file of the human proteome (UniProt KB, downloaded 7/16/2015, 68,554 entries) was added, and label-free quantification (LFQ) and matched between runs were checked. All other parameters were left as standard. The proteinGroups.txt file generated from the search was saved as a .csv, then statistics were calculated in R. The p-values are from an independent, two-sample t-test. Volcano plots were produced with the EnhancedVolcano R package.

**Plasmids**

Plasmids used in this study are as follows: pSAT1-RNF146-WWE-FLAG, pHis-TEV-AK2, pET-SUMO-CAPRIN1, pET-SUMO-G3BP2, pET-SUMO-DDX6, pET28a-PARP1. pSAT1 is a pBAT4-derived vector encoding a N-terminal 6x-HisSUMO tag. pHis-TEV-AK2 was purchased from Addgene (#38829). pET30-2-GAPDH was purchased from Addgene (#83910). pET-SUMO-CAPRIN1, pET-SUMO-G3BP2 and pET-SUMO-DDX6 were from J. Paul Taylor (St. Jude’s). pET28a-PARP1 was from John Pascal (Université de Montréal).

**Expression and purification of UHRF1**

Plasmid containing full-length UHRF1 (pEXP-CT) was transformed into *E. coli* strain T7 Express (New England Biolabs). Cells were grown at 37°C in LB media supplemented with ampicillin to an O.D. 600 between 0.5-0.7. At this point, protein expression was induced with 0.2 mM IPTG and the temperature reduced to 15°C. Cells were harvested after ~16 h. Pellets were resuspended in lysis buffer (500 mM NaCl, 20 mM Tris-HCl pH 8.0, 10 mM Imidazole pH 8.0 and 10% Glycerol) and lysed using an Emulsiflex-C5 homogenizer (Avestin). Clarified lysates were applied to IMAC nickel affinity resin (Bio-Rad). Resin were washed and bound protein eluted with lysis buffer supplemented with 250 mM Imidazole. The sample was then applied to a HiLoad 26/600 S200 Superdex column (GE Healthcare) equilibrated with running buffer (20 mM Tris-HCl pH 8.0 and 250 mM NaCl ). Protein purity was confirmed by SDS-PAGE.

**Expression and purification of RNF146-WWE, AK2, CAPRIN1, G3BP1/2 GAPDH and DDX6**

Proteins were expressed in *Escherichia coli* strain DE3 Rosetta. PARP1 was expressed and purified according to literature precedent.[4] For RNF146, AK2, CAPRIN1, G3BP2, GAPDH and DDX6, transformed cells were grown in LB supplemented with the appropriate antibiotics at 37 °C / 200 rpm to an $OD_{600}$ of ~0.8 , cells were chilled on ice for 1 h, induced with 0.3 mM IPTG, then expressed at 16 °C / 200 rpm for 16-20 h. Cells were harvested by spinning at 3,000 *g* for 30 min, then flash-frozen and stored at -80 °C until purification. All purification steps were performed on ice or at 4 °C. Cell pellets were resuspended in lysis buffer (20 mM sodium phosphate pH 7.5, 300 mM NaCl, 25 mM imidazole, 0.5 mM TCEP, 5% glycerol) supplemented with 1X SIGMAFAST protease inhibitor cocktail, then lysed by sonication. For CAPRIN1, G3BP2, GAPDH and DDX6 the lysis buffer also contained 5 mg of RNase A

(ThermoFisher) per L culture. Lysate were clarified by spinning at 24,000 *g* for 30 min. Lysates were applied to HisTrap columns (Cytiva) or HisPur resin (ThermoFisher), washed with lysis buffer, then eluted in lysis buffer containing 300 mM imidazole. RNF146-WWE-FLAG was desalted into lysis buffer, then the 6xHisSUMO tag was removed by incubation with 6xHis-tagged SUMO endopeptidase at 4 °C overnight. The tag and protease were removed with a second round of Ni-NTA chromatography. Untagged RNF146-WWE-FLAG was further purified by gel filtration with a HiLoad 16/600 Superdex 200 pg (Cytiva) using gel filtration buffer 1 (20 mM Tris pH 7, 200 mM NaCl, 1 mM DTT, 5% glycerol). CAPRIN1 was further purified with a Q FF column (Cytiva) using low salt buffer (50 mM HEPES pH 7.5, 50 mM NaCl, 0.5 mM TCEP) and high-salt buffer (50 mM HEPES pH 7.5, 1 M NaCl, 0.5 mM TCEP). CAPRIN1, G3BP2, GAPDH and DDX6 were further purified by gel filtration with a HiLoad 16/600 Superdex 200 pg (Cytiva) using gel filtration buffer 2 (50 mM HEPES pH 7.5, 200 mM NaCl, 0.5 mM TCEP, 10% glycerol). All chromatography steps were performed on an NGC chromatography system (Bio-Rad).

**Biotin-PAR pull-down**

A mixture of PAR (2- to ~100-mer) was labeled with biotin using ELTA and purified with DHBB-agarose as described above. Protein (5 µM) was mixed with PAR (50 µM, monomeric concentration) and binding buffer (20 mM HEPES pH 8, 100 mM NaCl, 0.5 mM EDTA, 0.5% v/v NP-40) in a reaction with a total volume of (40 µL). Samples were incubated on ice for 1 hr. An aliquot (10 µL) was taken and mixed with SDS sample loading buffer as the input. Magentic streptavidin beads (10 µL, Pierce) were equilibrated with 1X binding buffer, then the samples were added to the beads and the mixtures were incubated with end-over-end rotation at 4 °C for 1 hr. The beads were washed once with binding buffer, then the protein was eluted with 1X SDS sample loading buffer supplemented with 20 mM biotin by incubating at 95 °C for 10 min.

Inputs and elutions were separated with SDS-PAGE, then detected with the SimplyBlue total protein stain (ThermoFisher).

**Cy5-PAR electromobility shift assay**

PAR (4-, 8- 16- or 32-mer) was labeled with Cy5 using ELTA and purified with DHBB-agarose and ion-pairing $C_{18}$-HPLC as described above. Cy5-PAR (10 nM) was mixed with protein at the indicated concentration in binding buffer (10 mM Tris pH 7.5, 100 mM KCl, 1 mM EDTA, 0.1 mM DTT, 0.01 mg/mL BSA, 0.01% w/v OrangeG, 5% v/v glycerol) and incubated at ambient temperature for 1 h. Samples were separated with a native 5% tris-acetate polyacrylamide gel (37.5:1 acrylamide:bis-acrylamide ratio) by applying 10 V per cm of gel in tris-acetate buffer (40 mM Tris, 2.5 mM EDTA, 20 mM acetate pH 7.8) for 45 min. Cy5 signal was detected with a Typhoon (Molecular Biosciences). PAR was quantified with ImageStudio (Li-Cor), then plotted in Prism 8 (GraphPad). Dissociation constants were calculated with a sigmoidal dose-response curve by measuring the $EC_{50}$ ($K_D$ ~ [protein] at which half of the PAR is bound).

**Protein Domain and Functional Enrichment Analysis**

The Pfam protein domain enrichments were calculated with the DAVID functional annotation tool (https://david.ncifcrf.gov/summary.jsp). The enrichments and *P*-values for Tri-RGG, WWE, and macrodomain were calculated manually with an R script that determines *P*-value with the EASE score, which is the same modified Fisher's exact test used by DAVID. g:Profiler (https://biit.cs.ut.ee/gprofiler/gost) was used for functional enrichment analysis of the 743 PAR hit genes (2 unique peptides identified in both replicates, PARprolink / control ratio > 2), resulting in a list of 1935 statistically significant enriched terms (Benjamini-Hochberg FDR < 0.05). The hit genes list was treated as an unordered query, and statistical tests were conducted within a statistical domain scope of only annotated genes, selecting terms with sizes between 2 and 500 genes, considering the GO molecular function, GO cellular component, GO biological process, KEGG, Reactome and WikiPathways data sources. The Ensembl ID with the most GO annotations were chosen for all ambiguous genes (ABCF2, CSNK1E, FKBP4, LUC7L2, MATR3, PDE8, TARDBP). The STRING network was produced with Cytoscape (v3.8.0) using a confidence cut-off of 0.8. Functional enrichment analysis was performed in Cytoscape and nodes were colored according to metabolic pathways (hsa01100), RNA binding (GO.0003723), chromosome organization (GO.0051276) and DNA repair (GO.0006281).

##### Isoelectric point and molecular weight analysis

Gene lists (see Supplementary Datafile) were converted from gene symbols to RefSeq peptide sequences in the FASTA format. The molecular weights and isoelectric points were then calculated with the IPC stand-alone Python script.[5] The isoelectric point value used is the average of the seventeen methods used by IPC for pI calculations. Violin plots were produced with Prism 8.0 (GraphPad).

##### Supporting information references


1. Spitale, R. C.; Flynn, R. A.; Zhang, Q. C.; Crisalli, P.; Lee, B.; Jung, J.; Kuchelmeister, H. Y.; Batista, P. J.; Torre, E. A.; Kool, E. T.; Chang, H. Y. Structural imprints in vivo decode RNA regulatory mechanisms. *Nature* **2015***, 519*, 486-490.
2. Tan, E. S., Krukenberg, K. A., and Mitchison, T. J. Large-scale preparation and characterization of poly(ADP-ribose) and defined length polymers. *Anal. Biochem.* **2012***, 428*, 126-136.
3. Ando, Y.; Elkayam, E.; McPherson, R. L.; Dasovich, M.; Cheng, S.-J.; Voorneveld, J.; Filippov, D. V.; Ong, S.-E.; Joshua-Tor, L.; Leung, A. K. L. ELTA: Enzymatic Labeling of Terminal ADP-Ribose. *Mol. Cell* **2019**, 73 (4), 845–856.e5.
4. Langelier, M. F.; Steffen, J. D.; Riccio, A. A.; McCauley, M.; Pascal, J. M. Purification of DNA Damage-dependent PARPs from E. coli for Structural and Biochemical Analysis. *Methods Mol. Biol.* **2017**, *1608,* 431-444.
5. Kozlowski, L. P. IPC – Isoelectric Point Calculator *Biology Direct* **2016**, *11*, 55.

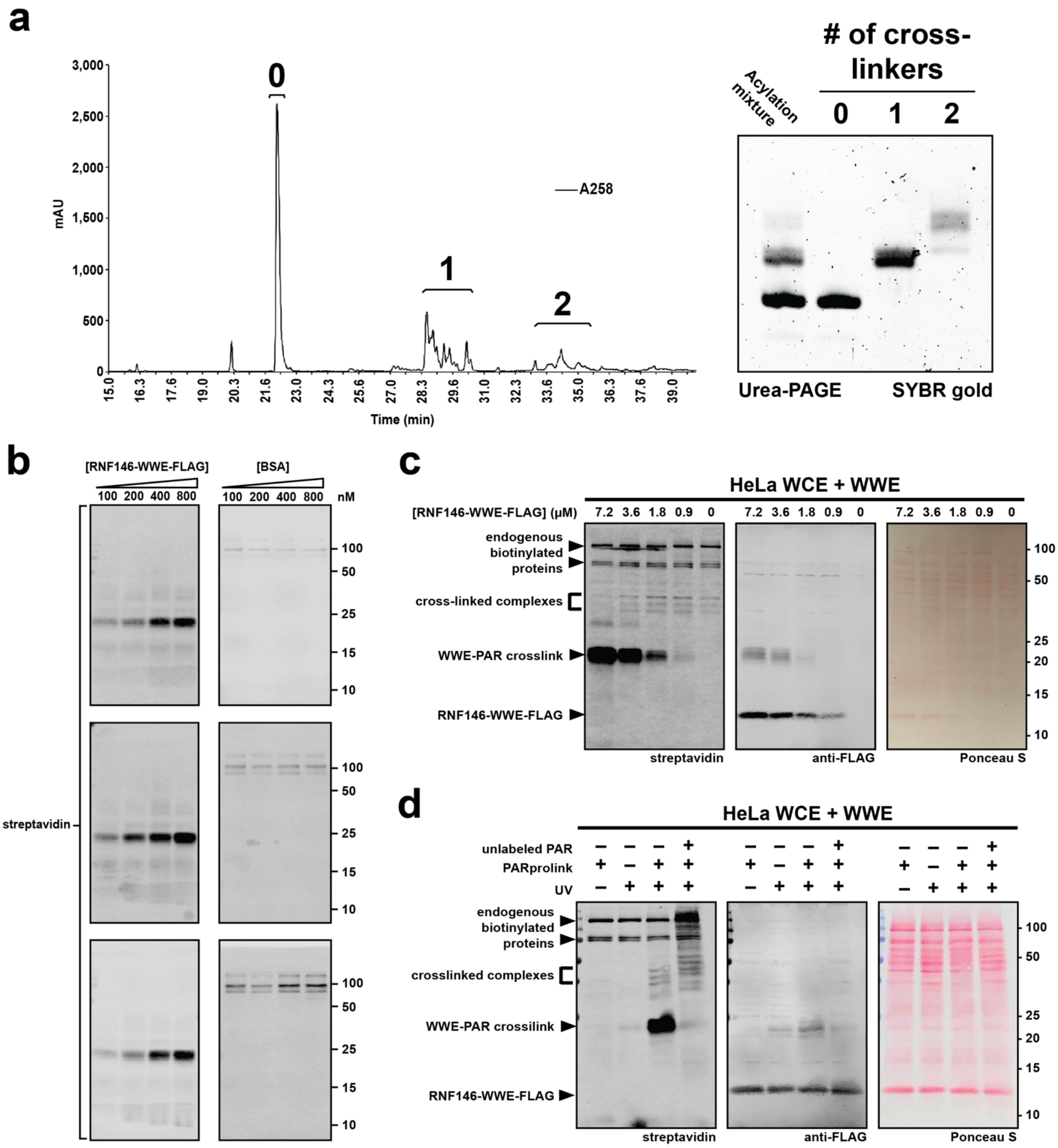


**Figure S1**. PAR photoaffinity probes capture a PAR-binding domain. (a) $C_{18}$-reverse phase HPLC UV chromatogram of a biotin-PAR mixture composed mostly of 0, 1 and 2 nicotinic acid analogues. Fractions corresponding to peak 1 were combined and used in photo-cross-linking experiments. (b) Full western blots from Figure 1c. (c & d) Photo-cross-linking recombinant RNF146-WWE-FLAG in HeLa whole-cell extract.

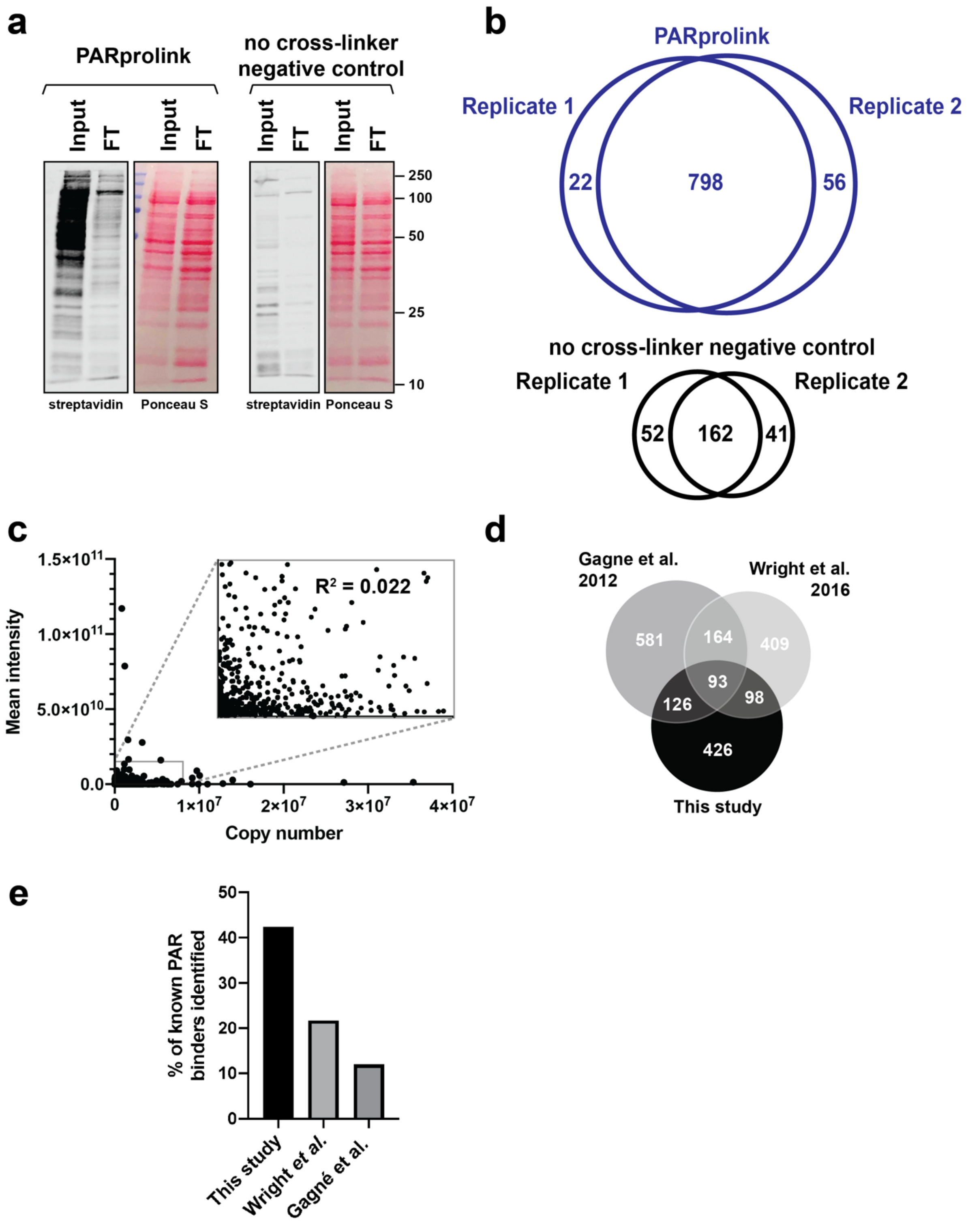


**Figure S2**. Photo-affinity-based isolation of the PAR-binding proteome in HeLa nuclear extract. (a) Representative western blots of the inputs and flow throughs from the samples submitted for LC-MS/MS. "Input" is the sample after UV cross-linking, but before addition of streptavidin agarose. "FT" is the flow through (supernatant) after streptavidin pull-down. (b) Venn diagrams depicting the overlap between two replicates of the photoaffinity PAR and no cross-linker control pulldowns. (c) Scatter plot of the proteins identified in our study, depicting the correlation between HeLa protein copy numbers (from Hein et al., 2015) versus the mean intensity observed in the PARprolink replicates. The R-squared value was calculated by a linear regression fit in Prism 8.0. (d) Venn diagram depicting the overlap between the proteins identified in our study and previously reported PAR interactomes. (e) Percentage of the 92 known PAR binders identified in our study and the previous PAR interactome studies.

**a**

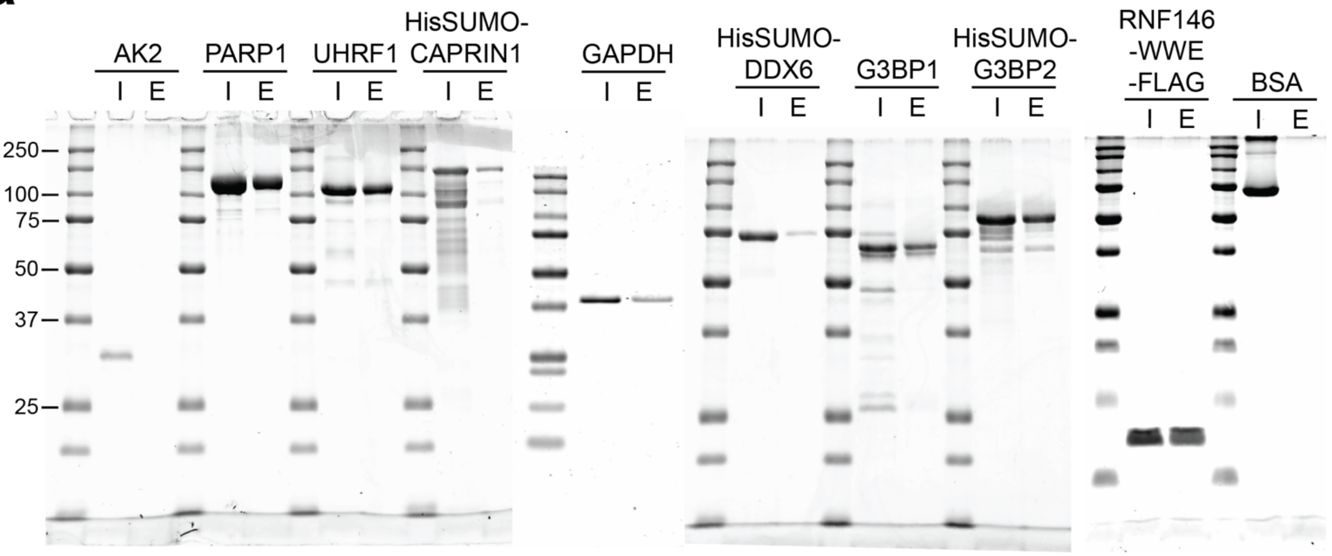


**b**

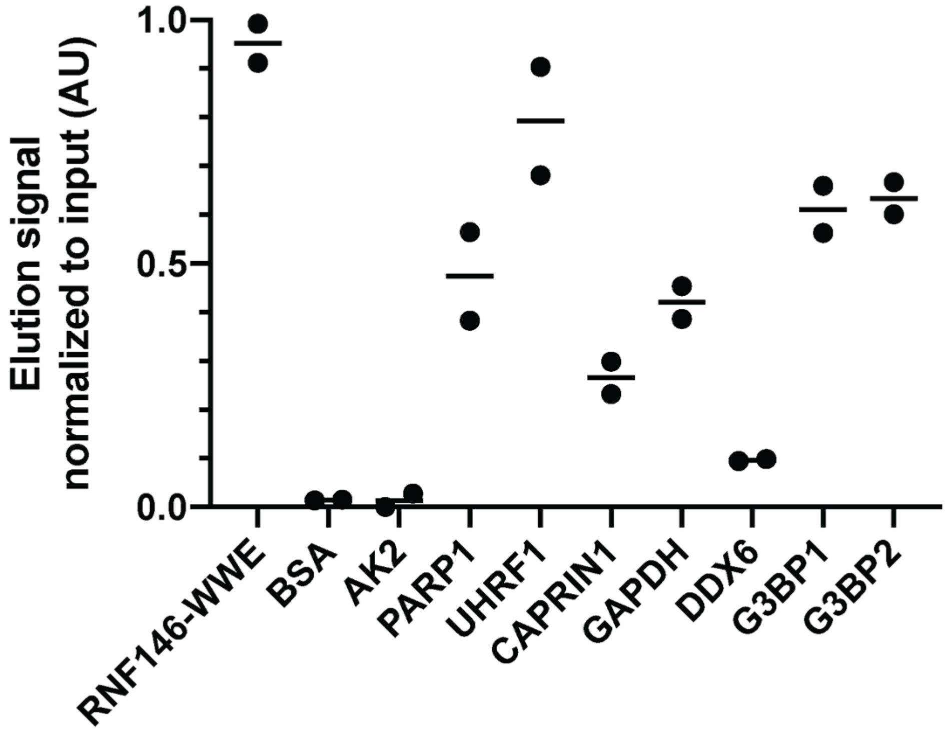


**Figure S3.** Biotin-PAR reverse pull-downs using a mixture of PAR lengths. (a) Uncropped gels from Figure 2c, I = input, E = elution. The data shown are representative of two independent experiments. (b) Quantification of biotin-PAR pulldown efficiencies from both replicates.

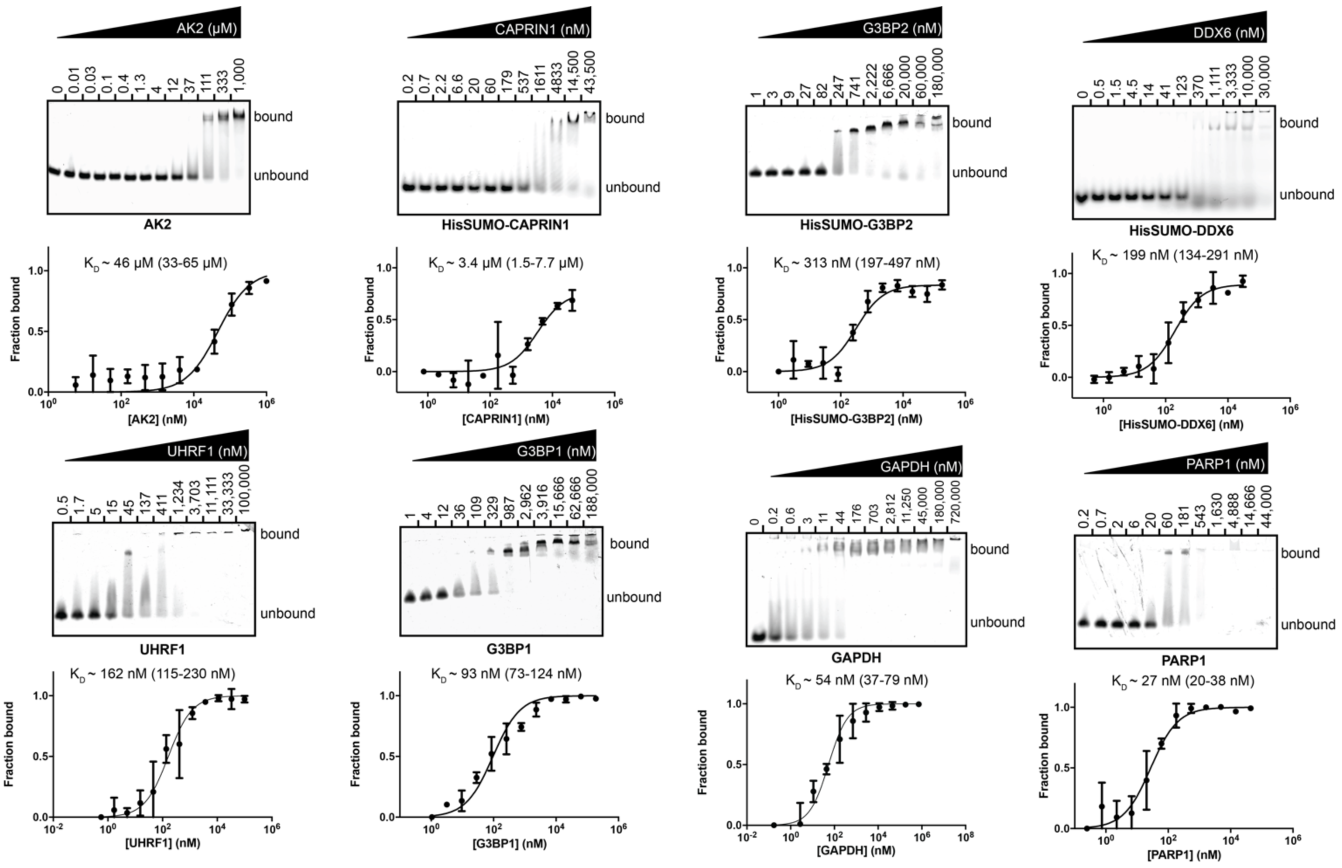


**Figure S4.** Representative gels from electrophoretic mobility shift assays (EMSAs) with eight PAR-binding candidates identified in the proteomics experiment. Data were quantified with ImageStudio and plotted with Prism 8 (values plotted are mean ± s.d., n = 3). To calculate dissociation constants, the plotted data were fit with a sigmoidal dose-response curve to calculate the $EC_{50}$ (~$K_D$ = [protein] at which half of the PAR is bound). Values in parentheses indicate the 95% confidence interval for the fit of the $EC_{50}$.

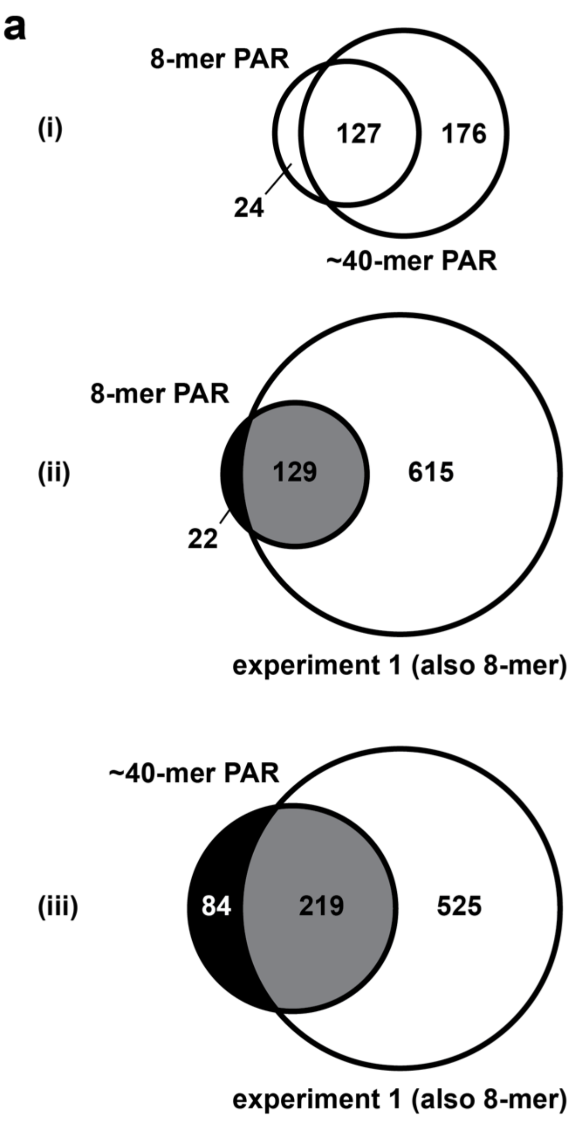


**b**

| PAR length | $K_D$ | 95% C.I. | $R^2$ | No. rep. |
|---|---|---|---|---|
| 4-mer | ~440 nM | 121–1570 nM | 0.71 | 3 |
| 8-mer | ~180 nM | 58–582 nM | 0.71 | 3 |
| 16-mer | ~27 nM | 11–68 nM | 0.82 | 3 |
| 32-mer | ~79 nM | 57–110 nM | 0.98 | 3 |

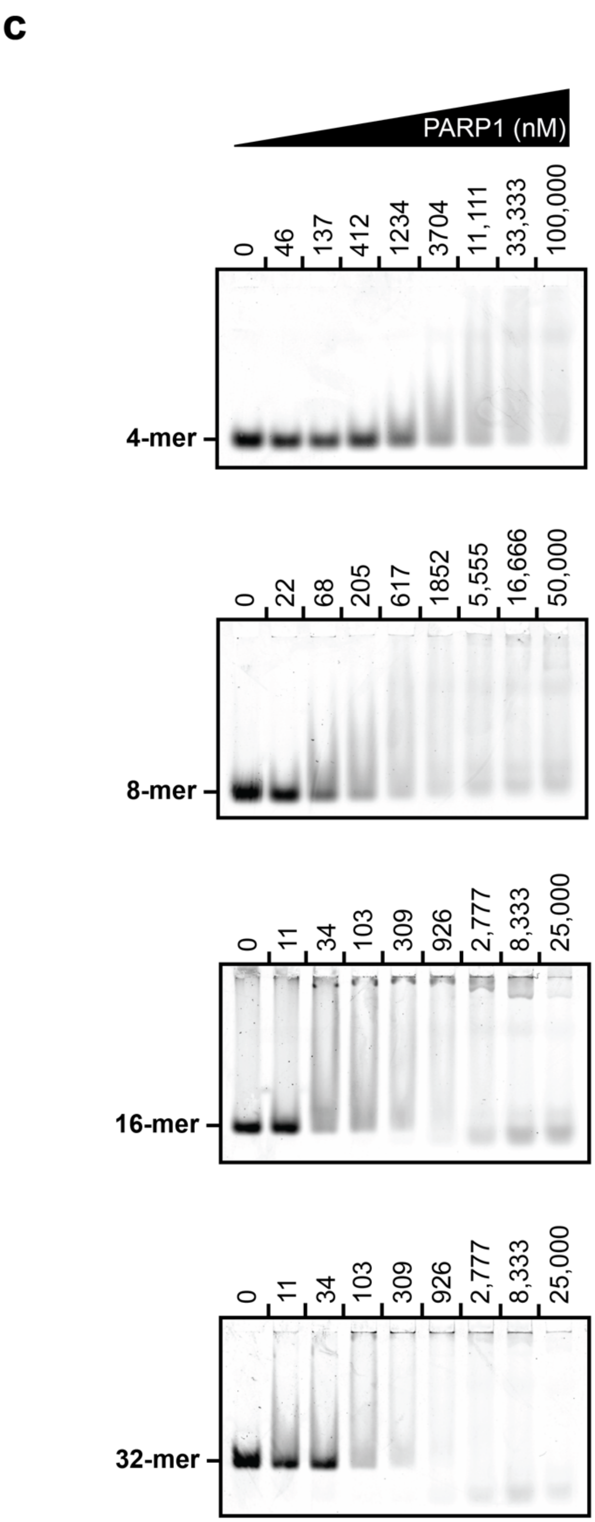


**Figure S5.** (a) Venn diagrams showing (i) the overlap between PAR-binding proteins identified in the 8-mer pulldown (151) and the ~40mer pull-down (303) (ii & iii) proteins from the 8mer and ~40mer PAR pull-downs compared to proteins identified in independent 8-mer experiments shown in Fig. 2. (b) Summary of the binding assay data from EMSAs with PARP1 and four different lengths of PAR. (c) Representative gels from the data summarized in panel b.

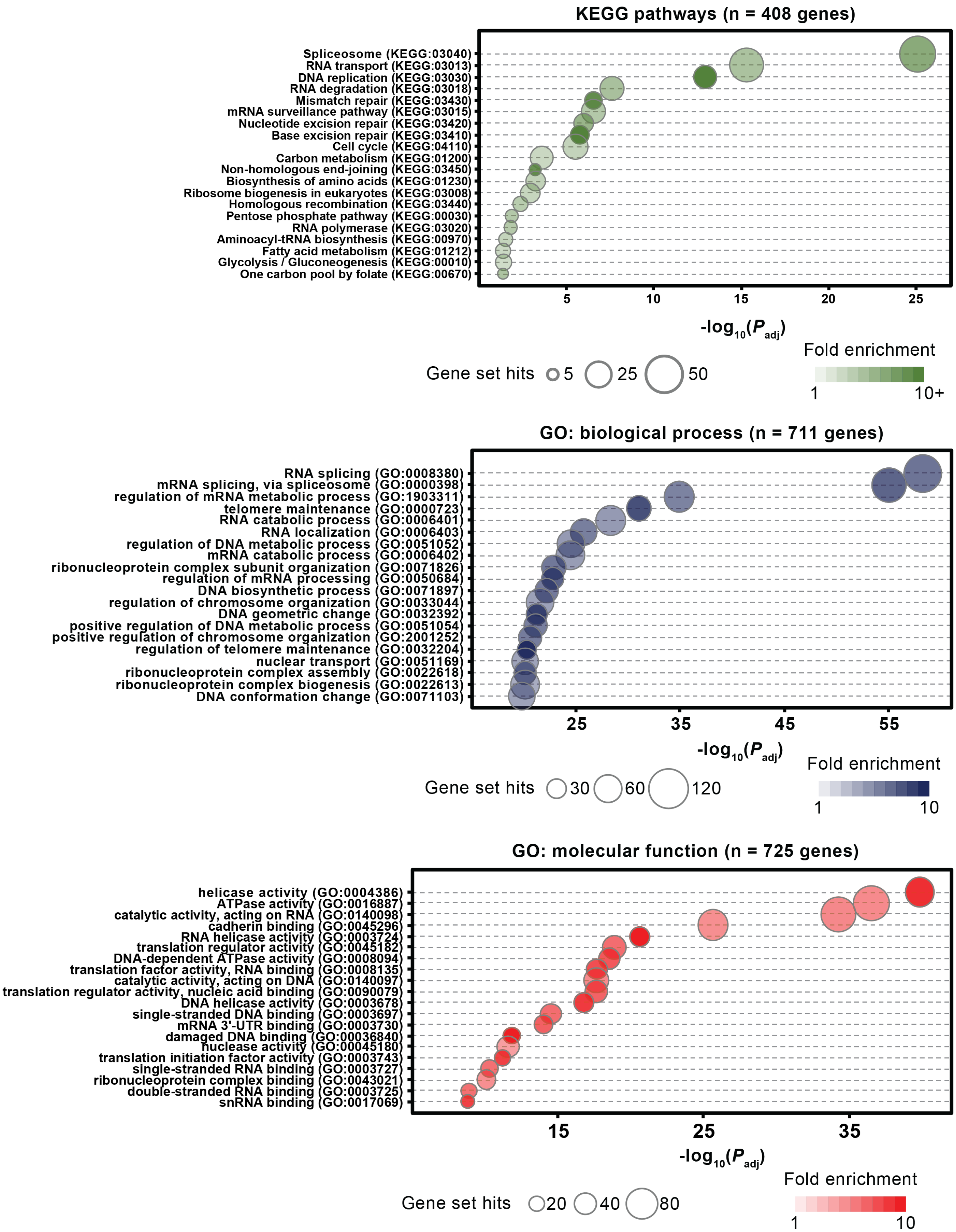


**Figure S6.** Top 20 enriched gene ontology terms for KEGG pathways, biological processes and molecular functions identified using g:Profiler (P < 0.05, with Benjamini-Hochberg FDR correction) among all 743 PAR-binding candidates.

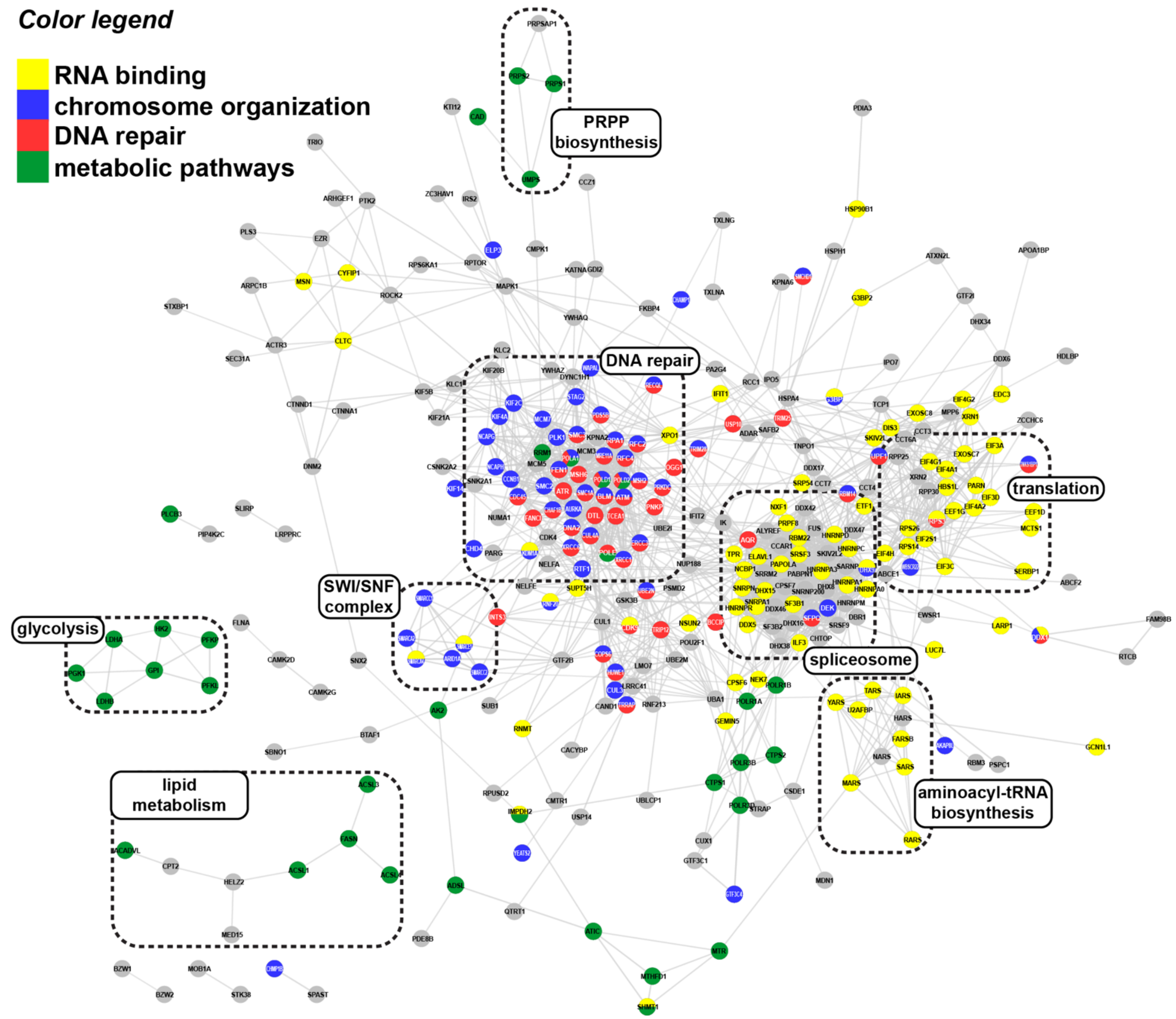


**Figure S7.** Network of high-confidence PAR-binding candidates (enrichment ratio > 8, P < 0.05, 416 genes) with genes annotated.

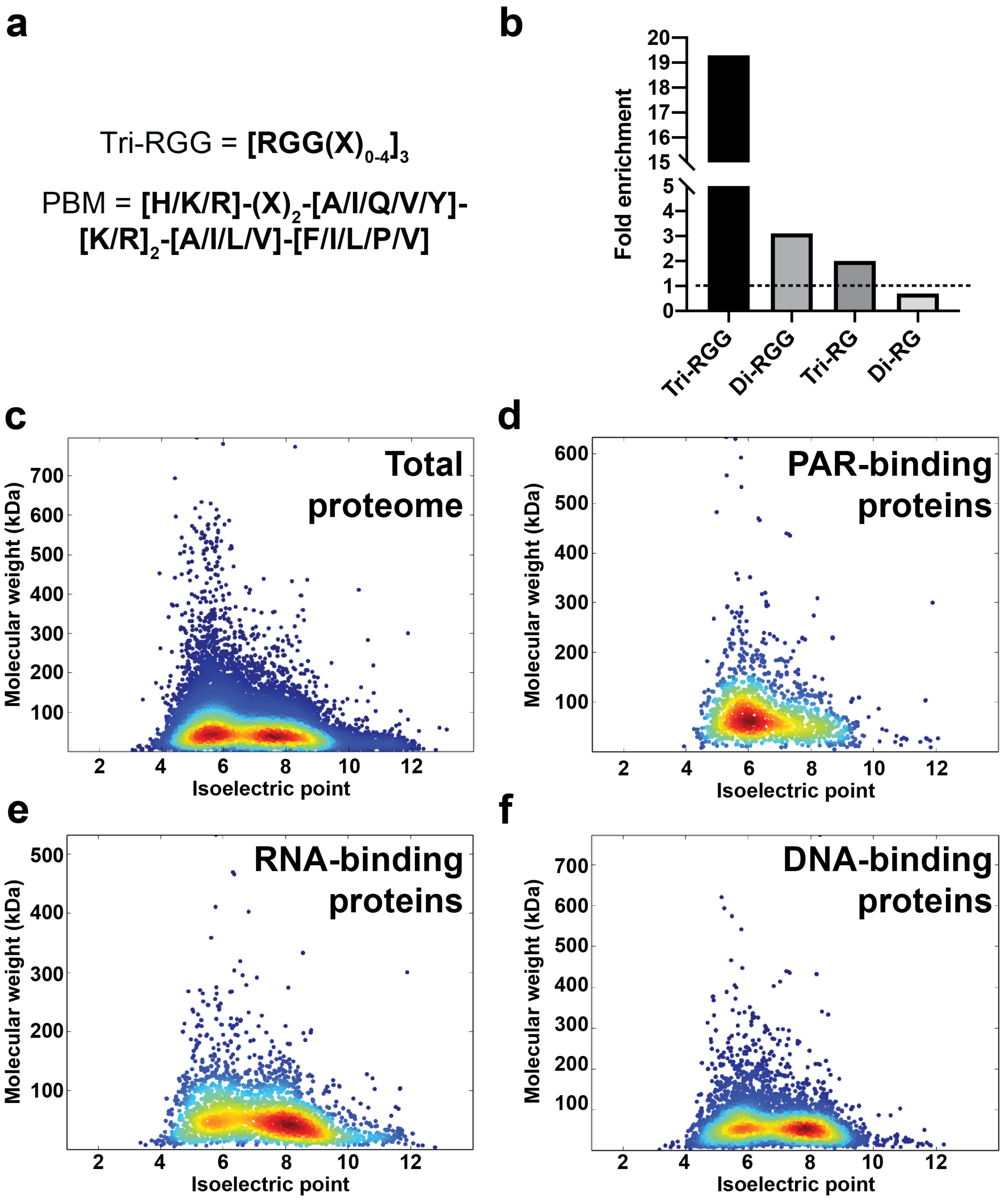


**Figure S8.** (a) PAR-binding sequence motifs reported in previous literature (Tri-RGG, Altmeyer et al., 2015; PBM, Gagné et al., 2008). (b) Fold-enrichment of the Tri-RGG motif compared to other RG/G motifs, (c-f) Molecular weights and isoelectric points for the whole proteome, PAR-binding proteins (the 743 hits from this study), RNA- and DNA-binding proteins (see Supplementary Datafile). Proteins with MW > 800 kDa were excluded from the plots.

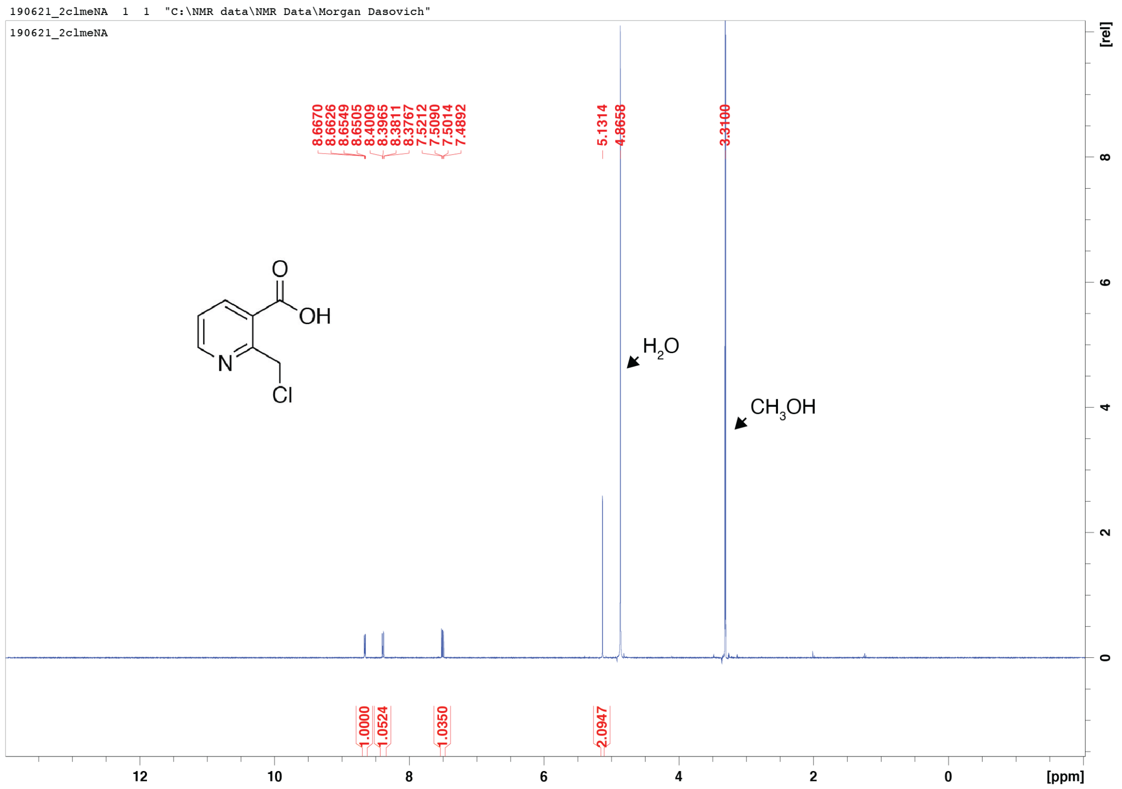


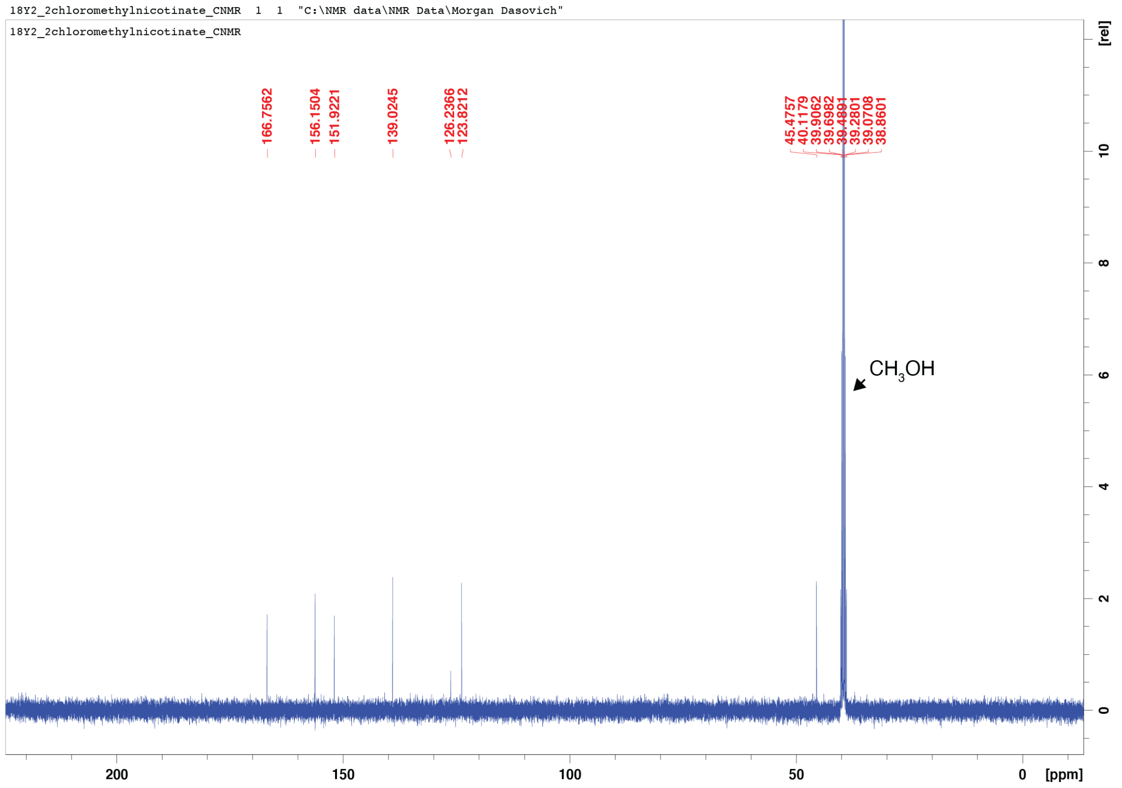


**Figure S9.** $^{1}H$ NMR (400 MHz) and $^{13}C$ NMR (101 MHz) spectra of 2-Chloromethylnicotinic acid ($CD_3OD$).

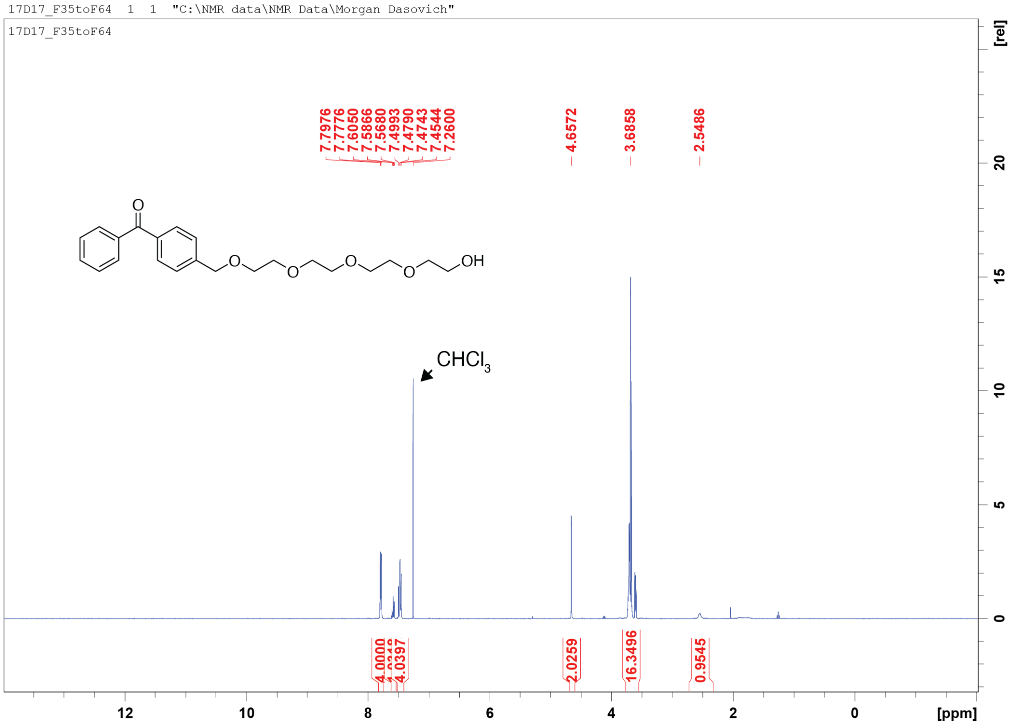


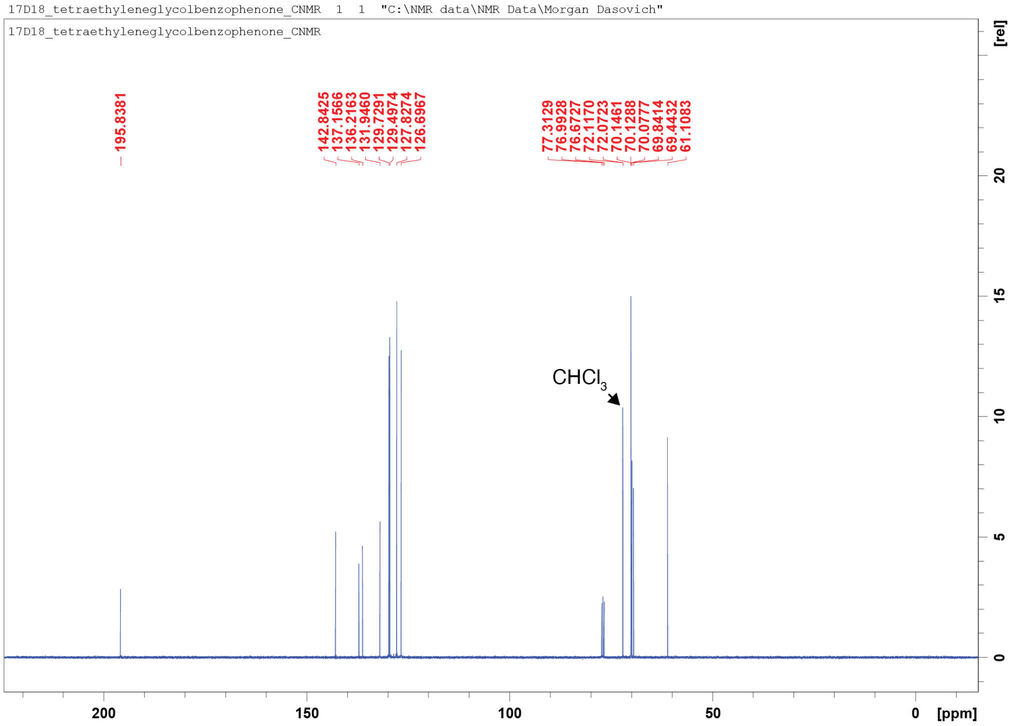


**Figure S10.** $^{1}H$ NMR (400 MHz) and $^{13}C$ NMR (101 MHz) spectra of (4-(13-Hydroxy-2,5,8,11-tetraoxatridecyl)phenyl)(phenyl)methanone ($CDCl_3$).

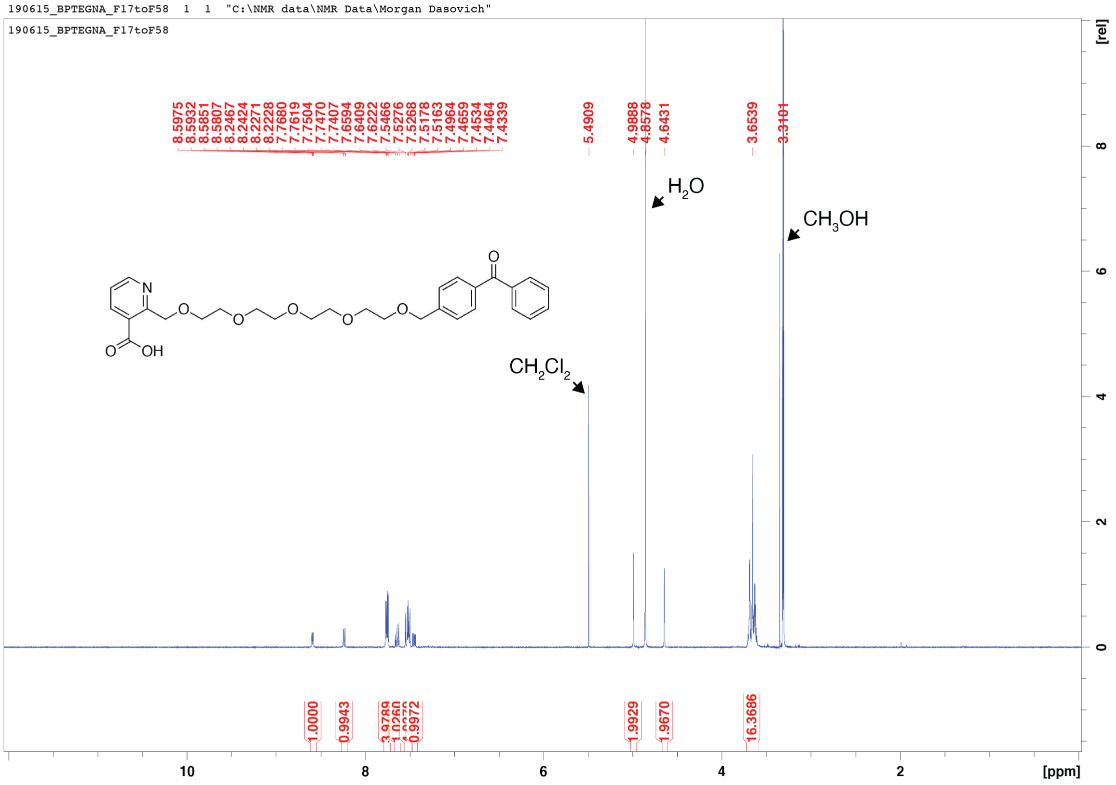


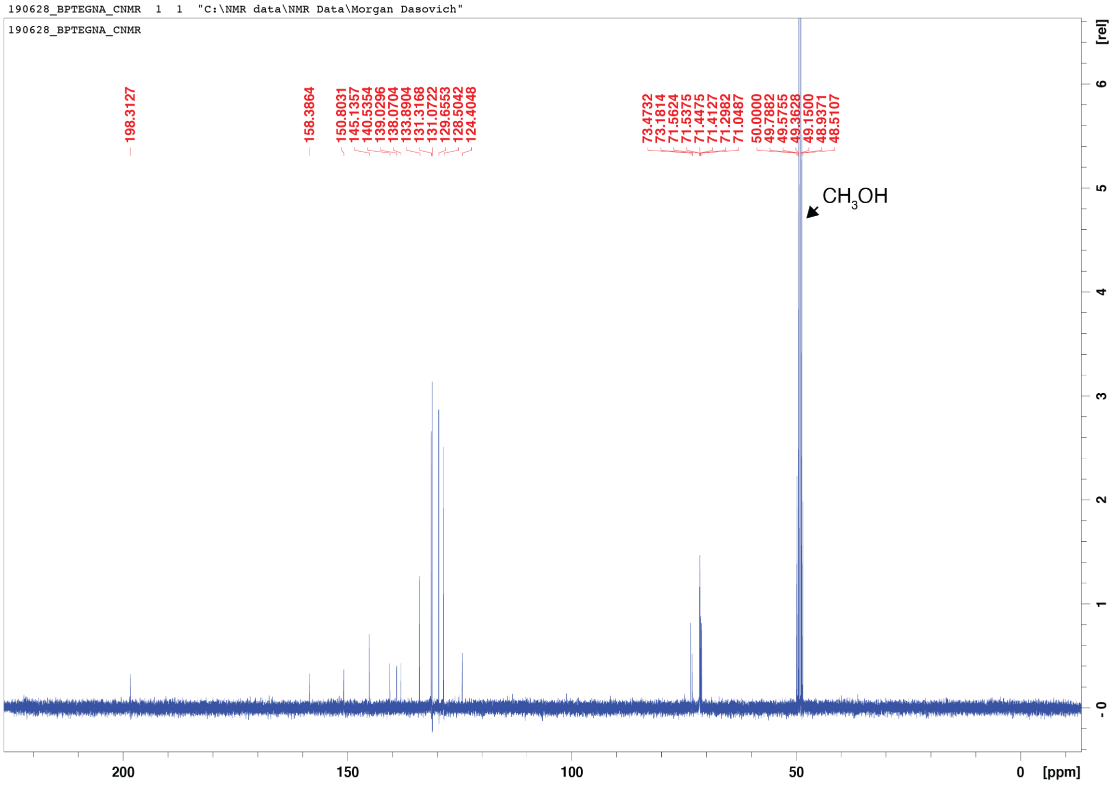


**Figure S11.** $^{1}H$ NMR (400 MHz) and $^{13}C$ NMR (101 MHz) spectra of 2-(15-(4-Benzoylphenyl)-2,5,8,11,14-pentaoxapentadecyl)nicotinic acid ($CD_3OD$).

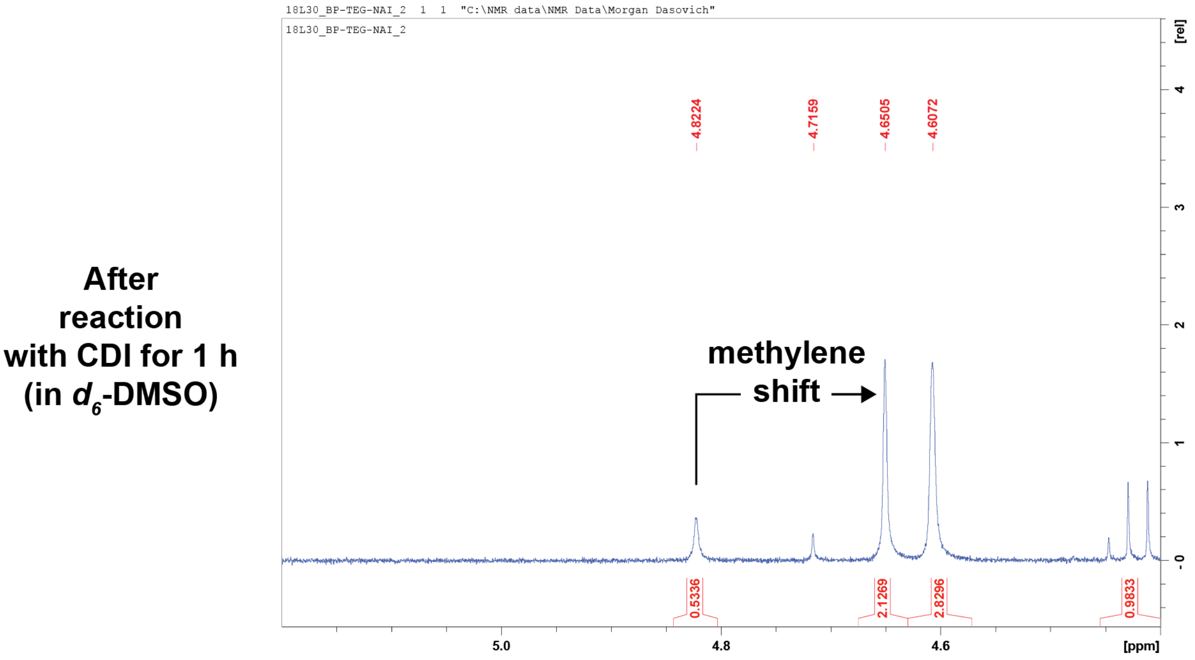


**Figure S12.** $^1$H NMR (400 MHz) of the reaction between 2-(15-(4-Benzoylphenyl)-2,5,8,11,14-pentaoxapentadecyl)nicotinic acid and carbonyldiimidazole ($d_6$-DMSO).
